# Uncovering the Signatures of Cellular Senescence in the Human Dorsolateral Prefrontal Cortex

**DOI:** 10.1101/2025.02.19.639091

**Authors:** Nicholas Sloan, Jason Mares, Aidan Daly, Lilian Coie, Shaunice Grier, Natalie Barretto, Obadele Casel, Kristy Kang, Chris Jackson, Maria Pedersen, Shruti Khiste, Ben Fullerton, Joana Petrescu, Courteney Mattison, Colin Smith, Yousin Suh, Vilas Menon, Hemali Phatnani

## Abstract

Identifying senescent cells poses challenges due to their rarity, heterogeneity, and lack of a definitive marker. We performed Visium spatial transcriptomics (ST) and single nucleus RNA sequencing (snRNA-seq) on non-pathological human tissue to build a transcriptomic atlas of aging and senescence in the dorsolateral prefrontal cortex (dlPFC). We identified markers characteristic of aging dlPFC cortical layers and cell types. We also observed an increase in astrocyte abundance and decrease in somatostatin expressing inhibitory neurons. Overall, the senescence profile in the dlPFC was highly heterogeneous and heavily influenced by cell type identity and cortical layer. Combined unbiased analysis of ST and snRNA-seq datasets revealed gene expression modules encoding for communities of microglia and endothelial cells in the white matter and regional astrocytes programs that were strongly enriched with age and for senescence-related genes. These findings will help facilitate future studies exploring the function of senescent cell subpopulations in the aging brain.

## Introduction

The dorsolateral prefrontal cortex (dlPFC) and its complex connectivity are critical for maintaining executive functions such as working memory, planning and maintaining attention^1,2^. dlPFC function progressively declines with age, evidenced by worsened performance in tasks assessing key executive functions among older adults^3–7^. A potential driver of this decline is cellular senescence, a central hallmark of aging^8,9^. Senescence encompasses heterogeneous cellular states triggered by prolonged cellular stress resulting in the irreversible arrest of cell proliferation and permanent changes to cell function *in vivo*^10–14^. Two main tumor suppressive pathways establish and maintain senescence: one governed by p53 and p21, the other by p16 and pRB^15^. Both pathways converge on downstream inhibition of CDK4 and CDK6 kinases, preventing cell cycle progression from G1 to S phase. Senescence plays an essential role in development^16^, wound healing^17^, and tumor suppression^18,19^, but accumulation of senescent cells is implicated in maladaptive processes such as carcinogenesis^19,20^, neurodegenerative disease^21,22^, and other age-related disorders^23^. Thus, senescence is considered antagonistically pleiotropic, promoting fitness in young age but driving aging and age-related pathologies late in life^24^.

Senescent cells accumulate in aged tissues^25–27^ including the brain^28–30^. Further, whole body clearance of senescent cells, either pharmacologically using a senolytic drug treatment of Dasatinib and Quercetin (D+Q) or genetically using INK-ATTAC^31^ mice treated with AP20187, resulted in improved cognitive function and extended healthy lifespan in mice^32^. Senescent cells are shown to actively contribute to aging and age-related pathologies through their senescence-associated secretory phenotype (SASP)^33–35^. The SASP is composed of pro-inflammatory cytokines, chemokines, exosomes, growth factors, and extracellular matrix-degrading proteins that are released following the induction of growth arrest^33–38^. Much of the diverse roles senescent cells play throughout the body can be attributed to differences in SASP composition, as separate senescent cell subpopulations have been theorized to promote tissue regeneration and inflammation^33,39^. The SASP has also been shown to have paracrine activities, inducing senescence in nearby cells^33^.

Post-mitotic cells, including neurons, are capable of expressing senescence-associated phenotypes *in vivo*^40^. In mice, there is an age-associated increase in neurons that display a senescence-like phenotype, including evidence of DNA damage, high ROS production, and p21 positivity^28^. In the human dlPFC, a senescence phenotype was observed in excitatory neurons exhibiting Alzheimer’s disease pathology, primarily marked by the presence of cell cycle arrest marker *CDKN2D*^41^, although the function of this marker in healthy neurons has not been fully explored. In addition to neurons, senescence markers have also been identified in microglia^42–44^, astrocytes^45–48^, and brain endothelial cells^49,50^. Microglia normally secrete many of the components considered to be part of the SASP, but increased secretion of TNF-α, IL1β, IL-6, and IL-8 have been observed in aged mouse microglia^51^. Senescent astrocytes positive for p16 have been shown to accumulate in the frontal cortex of Alzheimer’s disease patient brains^52^. Single-cell RNA-seq studies have shown an increase in endothelial cells expressing senescence-related genes in aged mice^53^.

Much of the evidence supporting the existence of senescent cells within the healthy aged human brain has measured the abundance of one or a few senescence markers at a time, and full scale transcriptomic or proteomic profiles of such populations have yet to be established. Since no single marker is distinctive of senescence, the current recommendation of the NIH Common Fund’s Senescence Network (SenNet) is to identify senescent cells by the presence of multiple known hallmarks at once^54^. Among these 9 proposed hallmarks are indicators of cell cycle arrest, the SASP, resistance to apoptosis, and the DNA damage response (DDR) pathway^54^.

The need to identify multiple markers to confidently identify senescent cells emphasizes the need for high-plex, high throughput technologies in assessing senescence in aged tissues. However, delineating which cells become senescent and identifying cell-type specific senescence marker panels remains a challenge. The identification of these heterogeneous molecular phenotypes has been hindered by our inability to collect *in situ* ‘omics’ data with sufficient depth and throughput to locate the small percentage of cells that exhibit a senescent phenotype. This challenge is compounded by a scarcity of appropriate study cohorts, including the availability of a broad age range of non-pathological human brain tissue. Here, we combine spatial and single nucleus genomics tools at scale, over multiple tissue sections from multiple subjects, at a depth sufficient to permit identification of the rare and heterogeneous senescence cell population. We apply these tools to non-pathological dlPFC tissue from the Sudden Death Brain Bank at the University of Edinburgh to generate an atlas of senescence in the human brain across the lifespan.

## Results

### Building a transcriptomic map of senescence in the dlPFC across the adult human lifespan

To build a transcriptomic map of senescence, we performed Visium spatial transcriptomics^55,56^ (ST) and single nucleus RNA sequencing (snRNA-seq) on each subject tissue in our assembled cohort. These unbiased transcriptomic techniques were utilized to capture changes in the diverse array of marker genes implicated to be involved in senescence hallmarks^54^ with accurate spatial and single cell resolution, respectively. To identify age-associated transcriptomic changes, postmortem dlPFC tissue was collected from 24 subjects with ages ranging from 33 to 74 years at the Sudden Death Brain Bank at the University of Edinburgh (Methods, Figure 1A, Table S1). In our 24-subject cohort, both ST and snRNA-seq were performed on the same block of tissue for 20 subject tissues; 2 subject tissues have only ST data, and 2 subject tissues have only snRNA-seq data. In-depth characterization of tissue samples and clinical data ensured that subjects were cognitively normal and lacked disease pathology (Table S1).

**Figure 1.**
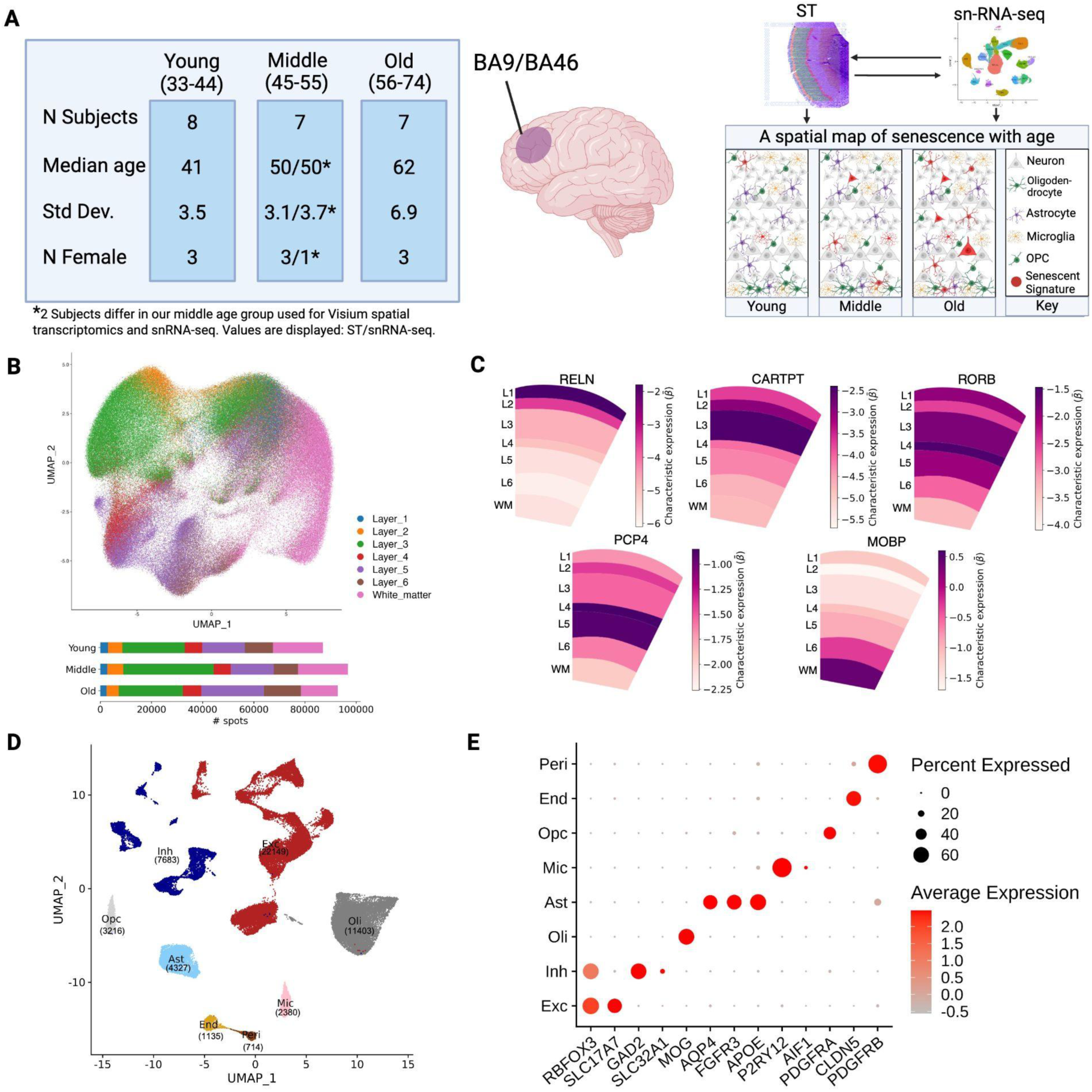
Building a transcriptomic map of senescence in the dlPFC across the adult human lifespan. **(A)** Summary table of subject cohort demographics for ST and snRNA-seq experiments (left). Diagram outlining the aims of this project (right). **(B)** UMAP embedding of ST data colored by cortical layer marker annotation (top). Bar plot depicting number of spots in each AAR label for young, middle and old subjects (bottom). **(C)** AAR level Splotch-predicted expression (*β*) – measured in log scaled counts of gene expression per spot within an AAR of interest (y-axis scale) – of key cortical layer markers within reference ST arrays from the middle aged group. L1-6 = Layer 1-6, WM = white matter. **(D)** UMAP embedding of snRNA-seq data colored by broad cell type class annotations. Number of nuclei profiles for each class displayed in parentheses. **(E)** Average normalized expression level of key cell type markers for annotated broad classes. Exc = excitatory neurons, Inh = inhibitory neurons, Oli = oligodendrocytes, Ast = astrocytes, Mic = microglia, Opc = Oligodendrocyte precursor cells, End = endothelial cells, Peri = pericytes.

### Identifying changes in the transcriptome with accurate spatial resolution with Visium ST

Using ST, we performed unbiased spatially-resolved transcriptome-wide sequencing on 80 tissue sections from 22 subject tissues profiled (Table S1). Spots within each Visium capture array were assigned to gray matter layers 1-6 and white matter. Anatomical annotation regions (AARs) were annotated based on the cell-type composition and density observed in our H&E stained tissues unique to each layer (Methods). After QC/filtering (Methods), our ST dataset reliably measured the expression of 13,702 genes across 276,701 ST spots. Dimensionality reduction of all ST spots in our dataset revealed that the majority of spot-level variability is attributable to our cortical layer annotation (Figure 1B), supporting the validity and use of these annotations. UMAP visualization additionally revealed minimal contribution of age grouping, sex, and Brodmann region to the overall variance in our dataset (Figure S1).

Since we expect that senescent cells will be sparse in non-pathological tissues, we used Splotch^57,58^, a hierarchical Bayesian statistical model of spatial gene expression, to integrate information across multiple tissue sections via our manual annotations in order to overcome the biological and technical sparsity of individual ST experiments. Using this model (Methods), we identified genes differentially expressed with age across cortical layers (Table S2). To validate whether our manual annotations replicated the known biology of cortical layers, we used Splotch to determine the expression of anatomical layer markers throughout our AARs. Importantly, we found that layer marker distributions within our AARs were in agreement with the biology described in the literature^59,60^, with *RELN* displaying the highest level of expression in annotated layer 1, *CARTPT* in layers 2 and 3, *RORB* in layer 4, *PCP4* in layers 4 and 5, and *MOBP* in the white matter (Figure 1C).

### Identifying changes in transcriptomic profiles with single-nucleus resolution using snRNA-seq

Visium ST generates accurate spatial maps of gene expression, but its resolution is limited by the size of each ST spot, which are 55uM in diameter with a 100uM center to center distance, and contain roughly 2-10 cells. Thus, to complement our spatial transcriptomic analysis, we used snRNA-seq to perform unbiased transcriptome-wide sequencing with single-nucleus resolution. Tissues used for snRNA-seq were sectioned in the same plane of the same tissue block as they were for ST experiments, in order to capture cell types within cortical layers represented in ST experiments. We profiled tissue blocks from 22 subjects and obtained transcriptomes of 53,007 nuclei after QC. Using unbiased clustering analysis, we identified 8 distinct clusters corresponding to the major cortical cell types (Figure 1D), including excitatory neurons, inhibitory neurons, oligodendrocytes, astrocytes, microglia, oligodendrocyte progenitor cells (OPCs), endothelial cells, and pericytes (Figure 1E).

### Cortical layer-specific age-associated markers

We used Splotch to determine differentially expressed genes (DEGs) between age groups within each cortical layer. We were interested in identifying both changes in gene expression conserved throughout the dlPFC, and changes within specific cortical layers. We determined the AAR-level characteristic expression pattern, (β), for each gene in each cortical layer for young, middle, and old age groups (Table S2). We used a Bayes Factor (BF) greater than 30 as a threshold to identify the most differentially expressed genes (Methods); these included 25 genes that were upregulated (Figure 2A) and 47 genes that were downregulated (Figure 2B) with age, comparing our young and old age groups.

**Figure 2.**
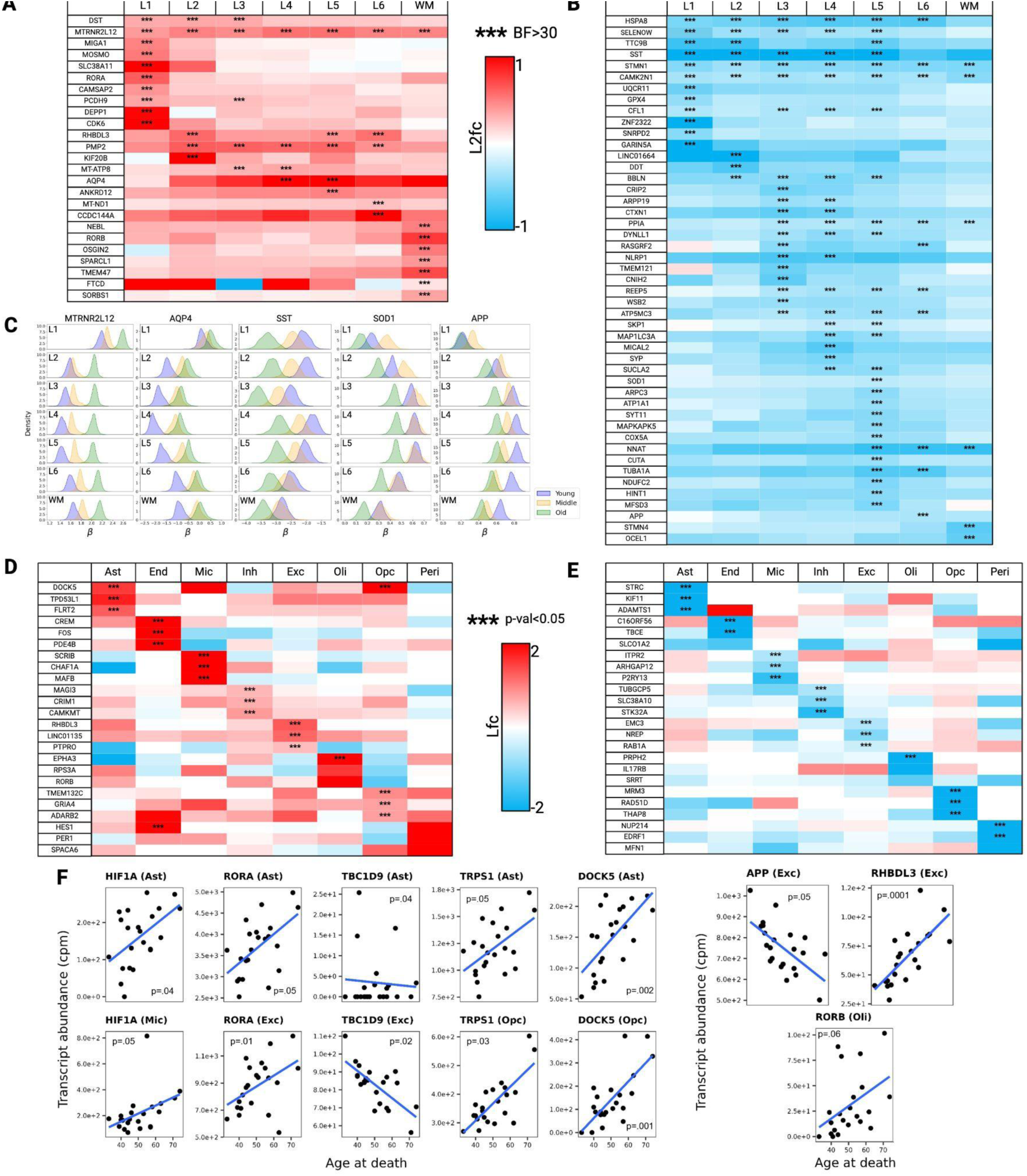
A spatial and single cell aging marker atlas of the human dlPFC. **(A-B)** Heatmaps displaying top ST differential markers across cortical layers between old (N=7) and young (N=8) cohorts. Top markers with higher expression in the old cohort in red shown in **(A)** and markers with lower expression in the older cohort in blue shown in **(B)**. Asterisks are added for markers with a BF > 30. **(C)** Kernel density estimate (KDE) plots displaying posterior distributions over expression (x-axis: log-scaled counts (*β*); y-axis: probability) inferred by Splotch for top aging markers across each layer (row) and age group (color code). **(D-E)** Heatmaps display top snRNA-seq differential markers across broad class cell types. Top markers with positive (red) and negative (blue) age association are shown in **(D)** and **(E)**, respectively. Asterisks are added for markers with a significant adjusted p-value (p<0.05). **(F)** Scatter plots highlight expression levels of top age-associated markers across age at death for specific broad cell class types. Post-hoc linear regression lines are added for clarity. Annotated p-value results are from differential expression analyses shown in Table S4.

Among the genes whose expression increased the most with age were *MTRNR2L12* and *AQP4.* While increased expression of *MTRNR2L12* was observed across all layers of the cortex, increases in *AQP4* were confined to layers 4 and 5 (Figures 2A and 2C). Genes whose expression decreased the most with age include *SST*, *SOD1*, and *APP* (Figures 2B-C). Decrease in *SST* expression with age was observed in gray matter layers 1-5, while decrease in *SOD1* and *APP* expression was confined to layers 5 and 6 respectively (Figures 2B-C).

### Cell-type specific aging markers determined using snRNA-seq

We also assessed gene expression changes with age in individual cell types. 452 genes were found to be changed with age (adjusted p-value <0.05) in various cell types, and almost half of these genes (221 of the 452) were differentially expressed within astrocytes (Table S4). Heatmaps summarizing top markers of broad class cell types increasing with age and decreasing with age are displayed in Figures 2D and E respectively. Age-associated trajectories of change in expression of a subset of these genes are shown in Figure 2F.

Interestingly, none of the age-related DEGs were shared by 3 or more cell types. 9 DEGs were shared by two cell types, among which 5 DEGs showed the same directional changes in expression (Table S4). We found that with age (Figure 2F): *HIF1a* increased in both microglia and astrocytes; *RORA* increased in both astrocytes and excitatory neurons; *TRPS1* and *DOCK5* increased in astrocytes and OPCs; and *TBC1D9* increased in astrocytes and excitatory neurons.

Moreover, analysis of the snRNA-seq and ST datasets together yielded converged predictions for age-related changes of certain genes. For example, in our ST data we observed that *APP* decreased with age in cortical layer 6 (Figure 2C) and snRNA-seq data showed *APP* decreased with age in excitatory neurons (Figure 2F). *RHBDL3* increased with age in cortical layers 2, 5, and 6 (Figure 2A), and *RHBDL3* also increased with age in excitatory neurons (Figure 2F). *RORB* increased with age in the white matter (Figure 2A), and *RORB* also increased with age nearing the threshold of significance in oligodendrocytes (Figure 2F).

### Change in the cellular composition of the aging dlPFC

In addition to changes in gene expression aggregated over AARs, alterations in cell type composition occur within the aged brain^61–63^. To investigate the dynamic changes in cell type composition during human dlPFC aging, we compared inferred cell type proportions within cortical layers across age. In addition to broad class cell types, we also explored age-related changes in cell type subclusters at a finer resolution by mapping our nuclei to a reference cortex dataset^64^ (Methods). After aggregating and merging some of the smaller subclusters, we were left with a total of 37 subclusters (Figure 3C).

**Figure 3.**
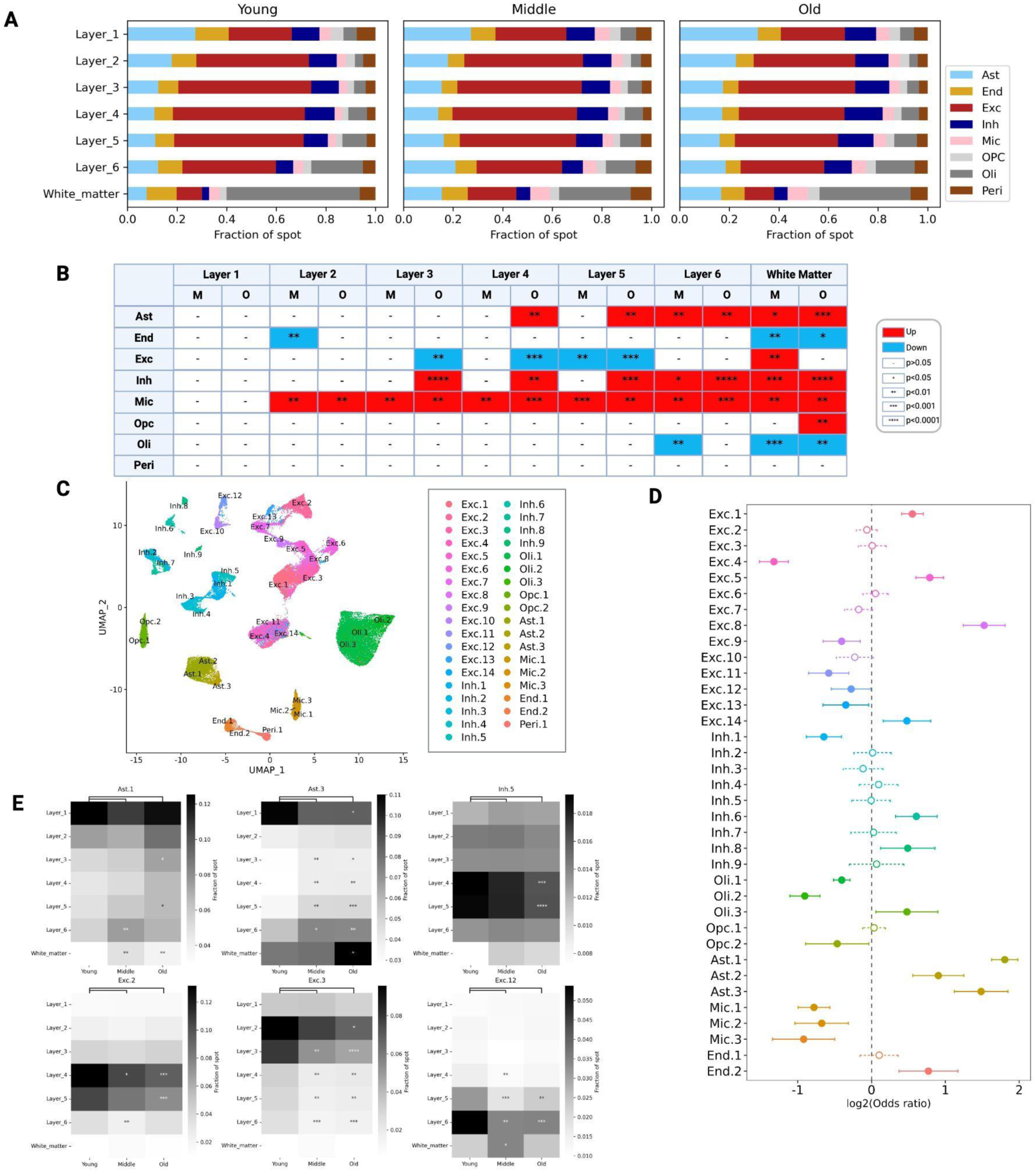
Changes in layer composition with age. **(A)** Cell2location cellular deconvolution of broad cell types across layers shown for the three age cohorts. **(B)** Table summarizes differential testing results of cell type composition between middle (M) or old (O) against young groups. Red (blue) cells indicate higher (lower) cell type enrichment of middle/old compared to the young group. **(C)** UMAP plot of subclusters of the snRNA-seq data. **(D)** MASC differential cell type proportion testing results between young and old groups are shown. Points to the left (right) indicate increased (decreased) cell type enrichment in the old group compared to the young group. Significant results (p<=.05) are indicated by solid, filled-in bubbles and solid confidence interval bars. **(E)** Heatmaps visualize abundance of six mapped clusters in ST data across cortical layers predicted by cell2location. Asterisks highlight significant results from differential abundance testing (BH-adjusted Wilcoxon ranked sum test) between middle and old, versus young groups. For heatmaps, asterisks indicate level of significance: ****=p<1e-4***=p<1e-3 **=p<1e-2, *=p<.05.

Synthesizing the ST and snRNA-seq datasets using cell2location^65^ – a deconvolution algorithm that predicts the abundance of cell types in multi-cellular spatial spots using a single-cell/nucleus reference – we found a significant increase in the inferred relative abundance of astrocytes with age in our ST data at the broad class level, especially in gray matter layers 4-6 and the white matter (Figures 3A-B). This layer-specific increase in astrocytes could primarily be attributed to subcluster Ast. 3, especially in layers 3-6 and the white matter (Figure 3E). Inferred proportions of subcluster Ast. 1 also increased with age in layers 3, 5 and the white matter. In addition, Mixed-effects modeling of Associations of Single Cells (MASC)^66^ analysis predicted a significant increase with age in the overall proportion of astrocytes in our single-nucleus data across all astrocyte subclusters (Figure 3D). Taken together, we found an overall increase in cortical astrocytes with age, particularly in the deeper gray layers of the cortex and the white matter.

We also investigated dlPFC neuronal populations potentially vulnerable to the aging process. The inferred relative proportion of excitatory neurons decreased with age, especially in gray matter layers 3-5 (Figures 3A-B). MASC analysis predicted a significant decrease of subclusters Exc. 4, 9, 11, 12 and 13 with age (Figure 3D), while cell2location predicted a significant decrease in Exc. 2, 3, and 12, in layers where these cell types were predominantly located (Figure 3E). Exc. 2 was mapped to layers 4 and 5, Exc. 3 to layers 2 and 3, and Exc. 12 to layers 5 and 6. In contrast, the inferred relative proportion of inhibitory neurons increased with age in layers 3-6 and the white matter (Figure 3B). However, at the subcluster level, we observed a significant decrease in Inh. 1 levels with age (Figure 3D), and a significant decrease in inferred Inh. 5 proportions in layers 4 and 5 (Figure 3E). As mentioned above, overall *SST* expression decreased with age in layers 1-5 (Figures 2B-C). Interestingly, both Inh. 1 and Inh. 5 express *SST* at a high level compared to the other Inh. subclusters (Figure S3). Taken together, these data suggest that there is a widespread change in the cellular composition of the aging brain, and highlight cellular subtypes that may be of particular interest in this regard.

### Cell types of the aging dlPFC vary in their expression of the canonical hallmarks of senescence

We next explored the expression patterns of known senescence hallmarks and curated gene sets across broad class cell types in our snRNA-seq dataset. This could potentially provide us with hints for cell type specific vulnerability to senescence inducing stressors, or cell type specific bias for certain senescent hallmarks. To accomplish this, we utilized SenMayo^67^ and Fridman Senescence Up^68^, two widely-used curated senescence gene lists, and a list of senescence target genes curated by the San Diego Senescence Tissue Mapping Center (SD TMC). We also incorporated a comprehensive list of p53-activated gene targets^69^, as p53 transcription factor activity drives a major growth arrest pathway that maintains senescence^15^. Finally, we utilized gene lists representing four of the nine senescence hallmarks proposed in the recent SenNet review paper^54^: Cell cycle arrest, SASP, DDR, and Resistance to Apoptosis. A summary of the genes present within each senescence hallmark list can be found in Table S9.

Each nucleus in our snRNA-seq dataset was assigned module scores quantifying the level of expression of each senescence hallmark gene list. Nuclei were deemed positive for a given hallmark if their module score ranked above the hallmark’s threshold (Methods). Overall, we observed major differences in the way that known senescence hallmarks present across dlPFC cell types. Our analysis predicted nuclei from excitatory neurons, astrocytes, and endothelial cells to be positive for a subset of the 8 mentioned hallmarks (Figure 4A). Microglia and pericytes have a significant enrichment of nuclei positive for all 8 senescence hallmarks (Figure 4A). Inhibitory neurons, oligodendrocytes and OPCs had an insignificant number of nuclei above threshold for all 8 hallmarks (Figure 4A). Additionally, we determined that microglia (12.5%), endothelial cells (13.5%), and pericytes (16.5%) displayed a significant proportion of nuclei positive for 3 or more senescence hallmarks at once (Table S10, Figure S5).

**Figure 4.**
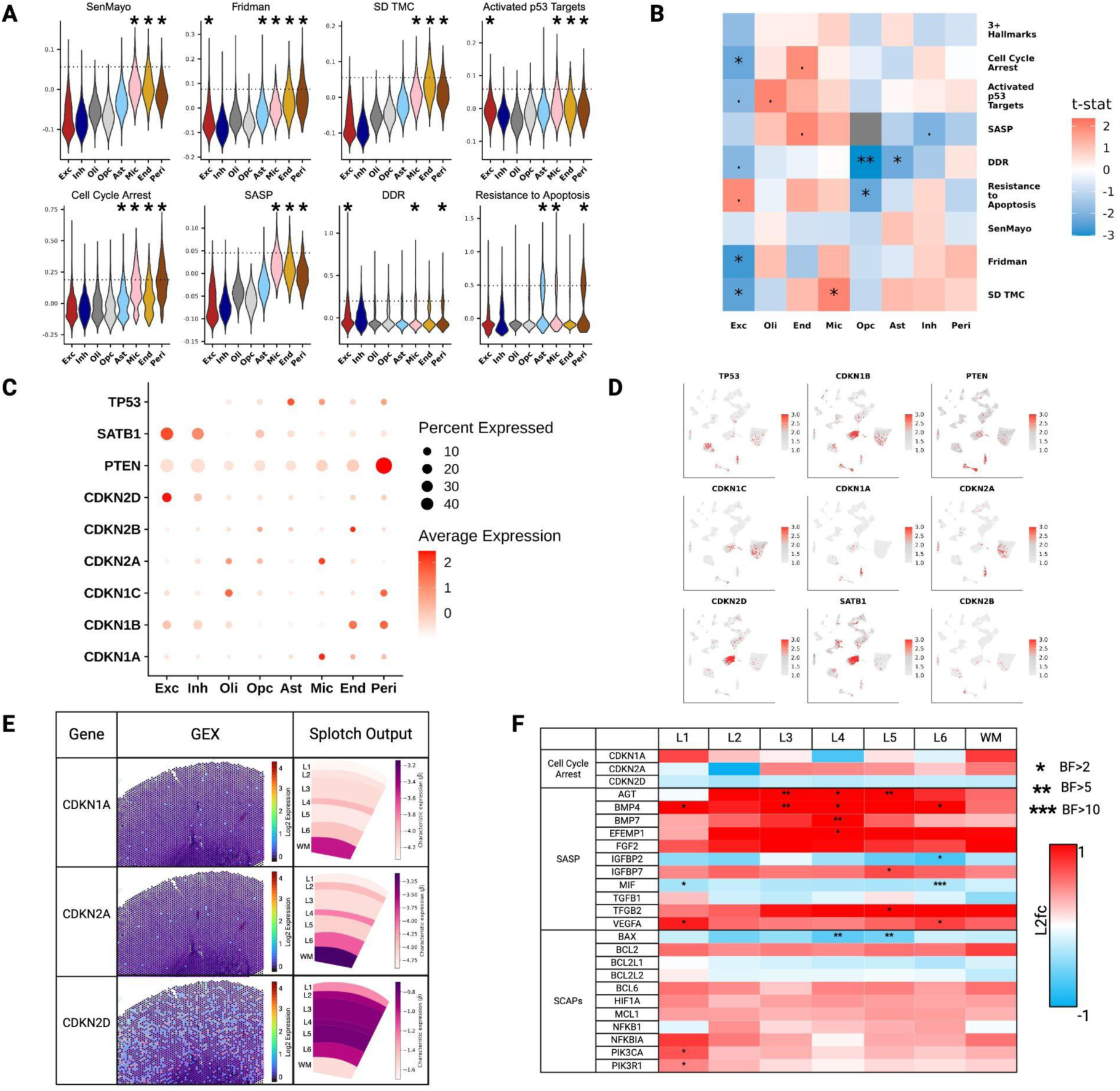
Survey of known senescence hallmarks across dlPFC cell types and cortical layers. **(A)** Distribution of module scores for eight senescence gene sets across broad class cell types are displayed. Each violin plot includes a dashed horizontal line indicating threshold for senescent positivity for specific hallmark. Stars are added to highlight statistically significant enrichment of positive senescence nuclei across broad class cell types. **(B)** Heatmap displaying enrichment of senescent-positive nuclei between young and old cohorts across senescence hallmarks. Cells represent test statistics from testing for difference of proportions via t-test. Red (blue) cells indicate an increased proportion of senescent nuclei in the old (young) cohort. Nominal p-values are annotated onto cells. **=p<1e-2, *=p<.05, .=p<=.10. **(C)** Average normalized expression levels of cell cycle arrest markers are shown, capping percentage expressed at 40% for better clarity. **(D)** Normalized expression patterns of cell cycle arrest markers are shown across the UMAP space of snRNA-seq data. **(E)** Grid showcases spot expression levels of key cell cycle arrest markers on a representative ST array (left) and Splotch predicted expression (*β*; color scale) of these markers across each annotated region (L1-WM) across all middle aged arrays (right). **(F)** Heatmap displays differential expression of genes associated with senescence hallmarks across cortical layers between young and old cohorts in our ST dataset. Increased (decreased) expression in the old cohort compared to the young cohort is shown in red (blue).

After determining which nuclei within the dlPFC display senescence hallmarks, we determined age-associated trends in the proportion of nuclei expressing senescence signatures. Of the cell types labeled positive for senescence hallmarks in Figure 4A, endothelial cells and microglia generally display positive trends in senescence hallmark expression with age, while excitatory neurons generally exhibit negative association of hallmark expression with age (Figure 4B). Of note, a significant increase in the number of microglia positive for the SD TMC’s senescence panel was observed with age, and an increase in the number of endothelial cells positive for cell cycle arrest and the SASP was trending towards significance with age (Figure 4B). Significantly fewer excitatory neurons were positive for Fridman Senescence Up with age, and a decrease in the number of excitatory neurons positive for activated p53 targets and DDR trended towards significance (Figure 4B).

### Cell cycle arrest marker expression is infrequent and heterogeneous throughout the dlPFC

We observed that certain dlPFC cell types were more likely to express specific cell cycle arrest markers. *CDKN1A* and *CDKN2A*, the genes encoding p21 and p16 respectively, were primarily expressed in subpopulations of microglia (Figures 4C-D), though expression of these two markers was seldom observed within the same cells (Figure S6). Additionally, *CDKN2D* and *SATB1* were primarily expressed in excitatory neurons, *PTEN* in pericytes, and *TP53* in astrocytes (Figures 4C-D). While astrocytes, which displayed *TP53* expression but not *CDKN1A* expression (Figures 4C-D), displayed an insignificant level of nuclei expressing the activated p53 target hallmark (Figure 4A), microglia, which displayed *CDKN1A* expression but not *TP53* expression (Figures 4C-D), were strongly associated with this hallmark (Figure 4A). These findings highlight the known disconnect between *TP53* mRNA levels and p53 transcription factor activity.

Similar to our snRNA-seq dataset, the canonical senescence markers are rare in our ST dataset. *CDKN1A* was expressed in 1.3% of ST spots and *CDKN2A* in 0.7% of ST spots (Figure 4E, Table S3). Importantly, for rare transcripts like *CDKN1A* and *CDKN2A*, it was impossible to discern a layer-specific pattern of expression from just one tissue section (Figure 4E), emphasizing the importance of multi-tissue integration via our Splotch analysis (Methods). Predicted cell cycle arrest marker expression patterns were consistent with the spatial pattern of cell types observed to express them, as *CDKN1A* and *CDKN2A* were primarily expressed in the white matter, while *CDKN2D* was expressed primarily in gray matter layers 3-5 (Figure 4E). Spot-level expression of *CDKN2D* was far more prevalent than expression of *CDKN1A* and *CDKN2A* (Figure 4E). This was unsurprising, as excitatory neurons are predicted to be far more prevalent than microglia across ST arrays (Figure 3A). Interestingly, we didn’t observe an age-associated significant difference in cell cycle arrest marker expression levels across cortical layers (Figures 4F, Table S2). This might be because expression of these transcripts is too rare to assess a trend with age across our subject cohort or that there are better markers that correlate more strongly with dlPFC senescence.

### Layer-specific enrichment of SASP factors in the aged dlPFC

In addition to assessing trends in cell cycle arrest marker expression with age in our ST dataset, we also quantified differential expression across known SASP factors^54,70–72^ and senescent cell anti-apoptotic pathway members^73,74^ (Figure 4F). We identified a layer-specific significant increase in SASP factors such as *AGT*, *BMP4*, *BMP7*, *EFEMP1*, *IGFBP7*, *TGFB2,* and *VEGFA* with age, especially in layers 3-5 (Figure 4F). *TGFB2*, *BMP4*, and *BMP7* ligands are each a part of the TGF-β superfamily and have been shown to play paracrine roles in the SASP, promoting cellular senescence in nearby cells^72^. In addition to increased expression of these ligands in the deeper layers of the aged cortex, we also saw a similar pattern of increased expression of key TGF-β family receptors *TGFBR3* and *BMPR1B* (Figure S6). *BMP4* has been shown to bind *BMPR1B* with high affinity^75^, and *TGFB2* has been shown to bind *TGFBR3* with high affinity^76^. Interestingly, all of the mentioned SASP factors and TGF-β family receptors that increase with age were expressed by astrocytes (except *BMP4*, Figure S6).

### Cell type-specific spatiotemporal dynamics of senescence-associated genes

We next investigated spatially coordinated patterns of senescence marker expression. We were interested in uncovering gene expression patterns encoding communities of senescent cells specific to subregions within the dlPFC. To identify changes in spatiotemporal patterns of gene expression with age, we applied a coexpression analysis to our spatial data (Methods). We identified 28 unique spatiotemporal gene expression modules, with each detectable gene in our ST dataset assigned to one module (Figure 5A, Table S5). Each of these modules were then subdivided into submodules based on the cell type-specific expression of each gene as assessed by our snRNA-seq dataset (Table S5). Each submodule can be thought of as a distinct cell type-specific expression program.

**Figure 5.**
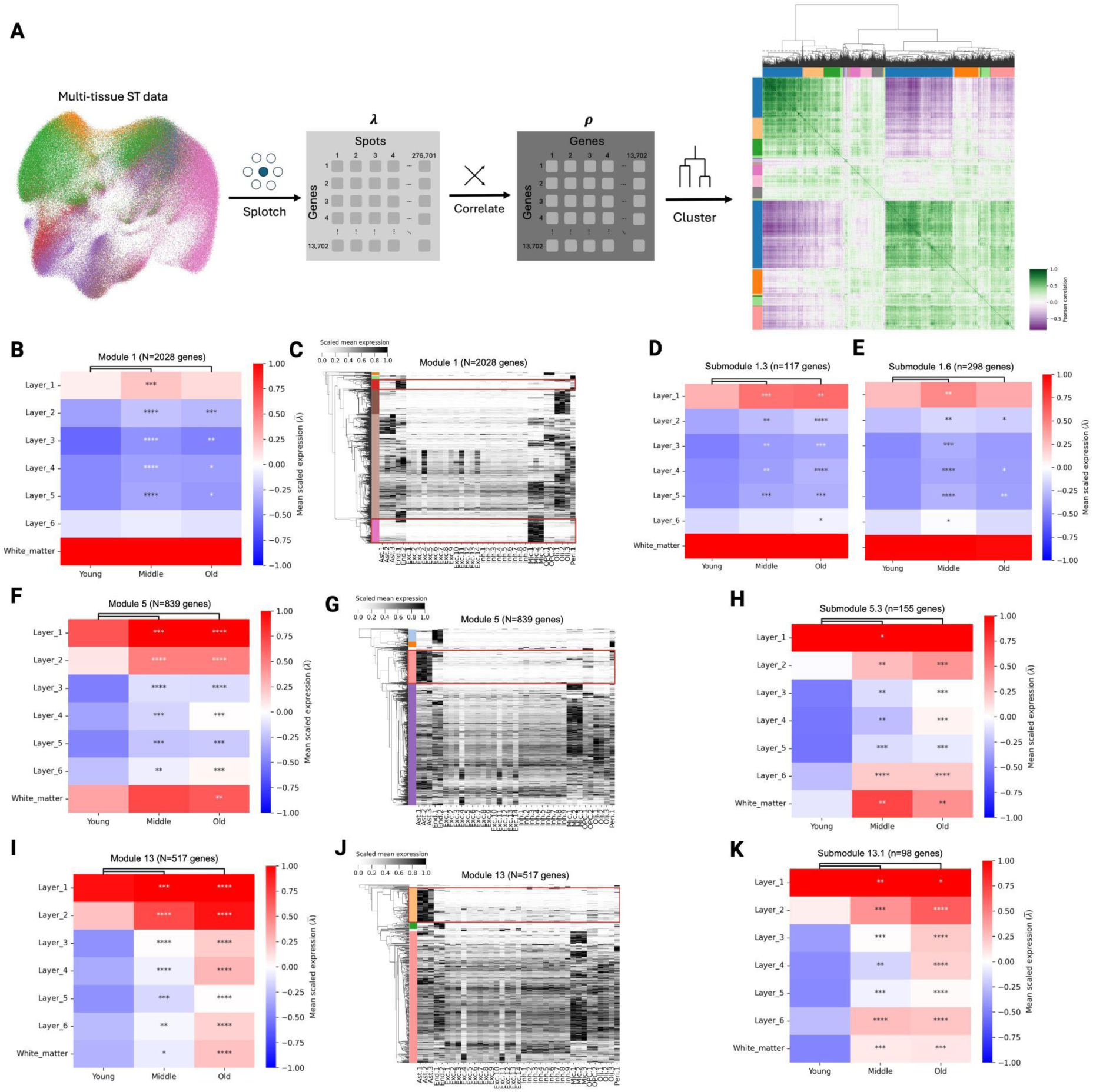
Spatial gene coexpression analysis reveals spatial coordinated cell type-specific senescence and age-related gene expression programs. **(A)** Unbiased pairwise gene co-expression modules generated as a function of spot level gene-gene covariance, yielding 28 unique modules. Pearson correlation is calculated for each gene pair comparing spot level expression values. **(B)** Expression level of Module 1 by cortical layer and age groups, with differential testing results between old and middle versus young. **(C)** Heatmap showcases expression patterns of genes in Module 1 across cell types using donor matched sn-RNA-seq data. Submodules 1.3 and 1.6 labeled with red boxes for emphasis. **(D-E)** Average expression levels of sub modules 1.3 and 1.6 across cortical layers and age groups shown, along with differential testing between middle/old and young groups. **(F)** Average expression levels of Module 5 by cortical layer and age groups. **(G)** Heatmap showcases expression patterns of genes in Module 5. Submodule 5.3 labeled with red box for emphasis. **(H)** Average expression levels of Submodule 5.3 across cortical layers and age groups shown. **(I)** Average expression levels of Module 13 by cortical layer and age groups. **(J)** Heatmap showcases expression patterns of genes in Module 13. Submodule 13.1 labeled with red box for emphasis. **(K)** Average expression levels of Submodule 13.1 across cortical layers and age groups shown. Asterisks indicate level of significance: ****=p<1e-4***=p<1e-3 **=p<1e-2, *=p<.05.

Among the 28 modules, Module 1 is enriched in the white matter (Figure 5B) and is significantly enriched for several gene ontology biological process (GO BP) pathways related to myelination and axon ensheathment (Table S6). Module 1 also contains a large proportion of the genes from the 8 senescence hallmark lists mentioned above (Figure S7). Gene expression within this module was primarily driven by non-neuronal cell types, such as endothelial cells, astrocytes, oligodendrocytes, and microglia (Figure 5C). For example, Submodule 1.6 is a microglia-specific submodule (Figure 5C) characterized by genes such as *AIF1*, *CD68*, *C1QA*, and *TYROBP* (Table S5). Submodule 1.6 is enriched in several GO BP terms related to inflammatory and immune response (Table S7), and contains most of the MHC class II genes in our dataset, such as *HLA-DRB1*, *HLA-DRB5*, *HLA-DMA*, *HLA-DRA*, *HLA-DPB1*, *HLA-DPA1*, and *HLA-DMB* (Table S5). This combination of markers and enriched GO processes led us to conclude that Submodule 1.6 encodes a population of reactive microglia in the white matter.

Submodule 1.6 displayed a significant enrichment with age, especially in the middle aged gray matter regions when compared to young subjects (Figure 5E). Submodule 1.6 also included senescence-related genes spanning multiple hallmarks, such as *CDKN1A, CDKN2A, TGFB1, TGFBR1, TGFBR2, TNFRSF1A, TNFRSF1B, SPP1,* and *ATM* (Table S5, Table S9). This result is consistent with our previous findings, as nuclei from microglia in our dataset were predicted to be positive for several senescence hallmarks (Figure 4A-B). Together, this potentially indicates a coordinated pattern of expression of senescence-associated genes within reactive microglia in the white matter of the dlPFC.

Submodule 1.3 is an endothelial cell-specific submodule that is enriched in endothelial cell type marker *PECAM1* as well as senescence markers *ATF3*, *B2M*, *ID3*, *IFI44*, *PLAT*, and *RRAS* (Table S5, Table S9). Submodule 1.3 expression significantly increased from young to middle to old across all gray matters layers (Figure 5D). GO BP pathway analysis revealed that the top enriched pathways within Submodule 1.3 were antigen processing and presentation of endogenous peptide antigen via MHC class I, endothelial cell differentiation, and negative regulation of cell population proliferation (Table S7). The majority of MHC class I genes in our dataset were expressed within this submodule, including *HLA-A*, *HLA-B*, *HLA-E* and *HLA-F,* in contrast to the MHC class II genes in Submodule 1.6 (Table S5). This result suggests that antigen-presenting cells share a similar spatial pattern within the dlPFC, primarily enriched in the white matter and cortical layer 1. Our snRNA-seq dataset similarly shows that MHC-I genes are almost exclusively expressed in endothelial cells, and MHC-II genes in microglia (Figure S6).

Exploring our spatiotemporal modules further, Modules 5 and 13 show a strong enrichment with age across all cortical layers (Figures 5F and 5I). Module 5 displays a spatial pattern favoring layers 1, 2 and the white matter, while Module 13 was mostly expressed in layers 1 and 2. These modules display distinct functional patterns – gene ontology analysis predicts Module 5 to be involved in tube morphogenesis, while Module 13 is more involved in chromatin organization (Table S6). Interestingly, both of these modules contain distinct astrocyte-specific Submodules, 5.3 and 13.1 respectively (Figures 5G, 5H, 5J, 5K).

Submodule 5.3 can primarily be identified by astrocyte cell type markers *SOX9* and *ALDH1L1*, known senescence markers *BCL2*, *DDIT4*, *IL33*, *FAS*, *FGF2*, *ORAI3* and *THBS4,* and TGF-β family receptors *BMPR1B* and *TGFBR3* (Table S5, Table S9). Submodule 13.1 contains the astrocyte gene marker *GLUL*, and known senescence markers *EGFR*, *F3*, *VEGFA*, and *TGFB2* (Table S5, Table S9). Thus, Submodules 5.3 and 13.1 represent distinct spatial subpopulations of astrocytes that are elevated with age and display different hallmarks of senescence.

### Neighborhood analysis of senescence modules with age

Our analysis thus far reveals layer-specific changes in expression patterns. To examine age-associated changes at higher spatial resolution, we focused on local, spot-level neighborhoods. We selected Submodules 1.3, 1.6, 5.3, and 13.1, eight curated senescence hallmark gene lists described above (Figures 4A-B), and publicly available GO and KEGG pathways representing processes known to be affected by senescence (Figure S10, Table S9). For each gene list, each ST spot in our dataset was assigned an expression score between 1 (lowest decile) and 10 (highest decile), and the “enrichment” – the propensity of spots with a given score to neighbor spots with another score – was calculated for all pairs of scores, separately for each age group (Methods). We were particularly interested in whether ST spots that scored highly for a particular senescence hallmark would tend to co-localize or would be more dispersed (Figure 6A-L). Additionally, we examined the distribution of spots per expression decile in each cortical layer (Figure S9). Generally, we would expect a skew towards higher expression of senescence modules with age. This can either result from an increase in high-scoring spots (d10), a decrease in low-scoring spots (d1), or both.

**Figure 6.**
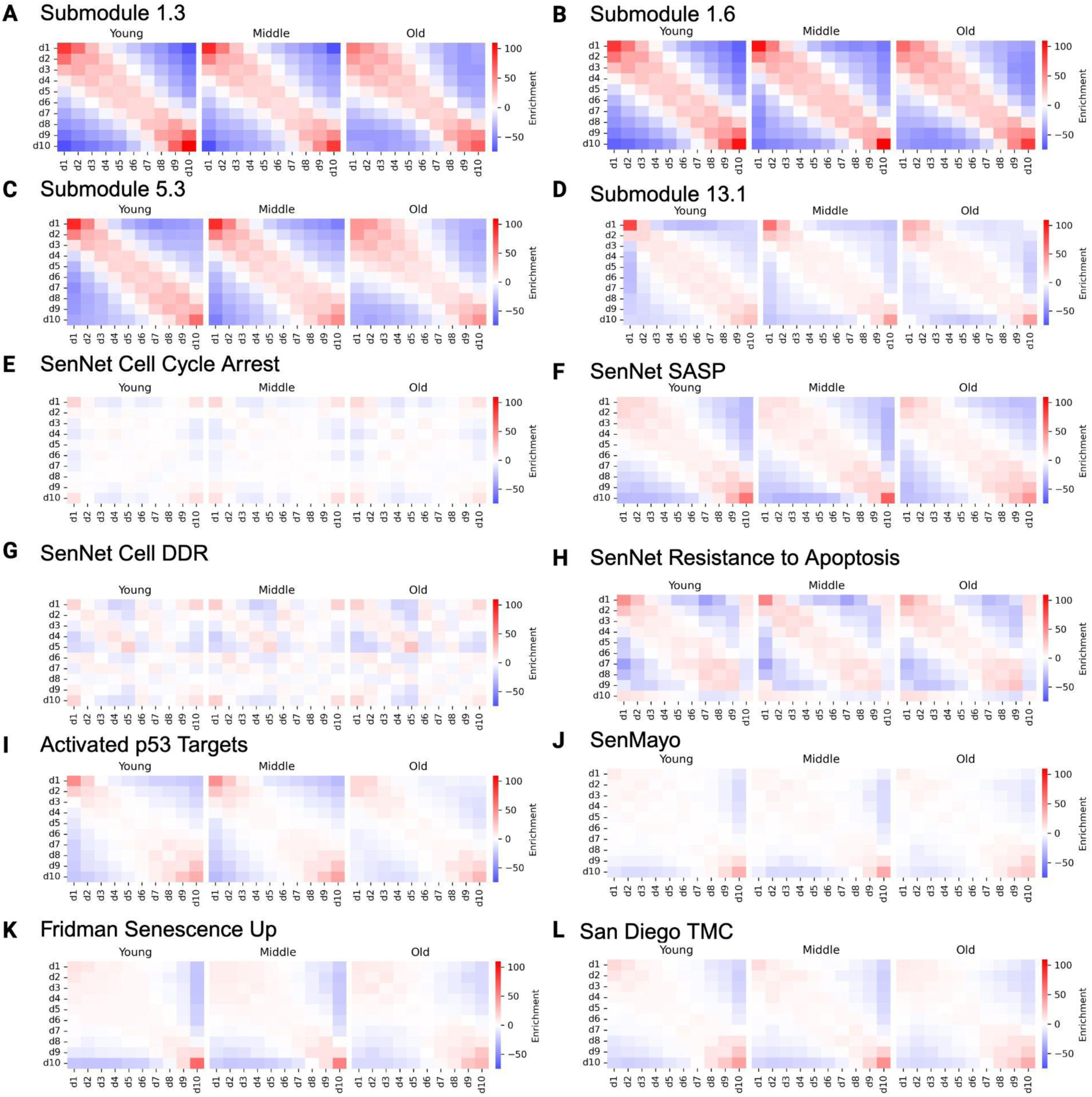
Spatial distribution of senescence modules with age. **(A-L)** Spatial transcriptomic spots were scored for expression of each unbiased spatial submodule of interest (**A-D**) as well as known senescence modules from the literature (**E-L**). Module scores across ST spots were divided into deciles. For each spatial submodule, we display heatmaps showing spatial enrichment statistics for each decile and quantified self-enrichment.

From our spatial enrichment analysis (Methods), most gene sets exhibited spatial coherence – spots with the same or similar expression score tended to neighbor each other. Visually, this is reflected by a strong diagonal signal on the pairwise enrichment plot (e.g., Figure 6A). Unsurprisingly, our spatially derived submodules (Submodules 1.3, 1.6, 5.3, and 13.1) showed a high degree of spatial coherence (Figures 6A-D). However, senescence hallmark lists curated from the literature, such as Senmayo, Fridman Senescence Up, and the SD TMC list, also displayed degrees of spatial coherence (Figures 6J-L). This indicates that spots predicted to be positive for senescence signatures tended to neighbor each other, potentially indicating that local external tissue signals in the dlPFC contribute to the induction of senescence phenotypes.

Exploring specific hallmarks of senescence also showed interesting results. While SASP signaling and resistance to apoptosis displayed a high degree of spatial coherence (Figure 6F, H), cell cycle arrest and DDR signals were more dispersed, with low-scoring spots often neighboring high-scoring spots (Figures 6E, 6G). SASP induction and resistance to apoptosis are often driven by external signals, such as paracrine SASP factors^77^. By contrast, cell cycle arrest and DDR induction in the dlPFC is perhaps driven by internal stimuli, causing spots positive for such signals to crop up in a seemingly dispersed manner. Alternatively, this result could reflect the high degree of heterogeneity and cell type specificity in cell cycle arrest marker presentation we observed across dlPFC (Figures 4C-D), and that cell cycle arrest in different cell populations occurs seemingly independent of one another. This is further supported by the fact that the p53 activation pathway, a specific pathway known to maintain cell cycle arrest, shows a high degree of spatial coherence (Figure 6I). Thus, this analysis provides use with hints about the potential mechanisms driving senescence hallmark induction.

## Discussion

Identifying senescent cells in non-pathological tissues poses challenges due to their rarity, heterogeneity, and the absence of a definitive marker. Evidence suggests that senescent cells accumulate in the brain with age^28–30^, yet the complex transcriptomic profiles of distinct senescent cell populations across cell types and brain regions remain unknown. In this study, we performed Visium ST and snRNA-seq on the same human tissue cohort to identify the signatures of aging and senescence in the dlPFC with both spatial and single-cell resolution.

We found that endothelial cells and microglia exhibited multiple known senescence hallmarks (Figure 4A) and that the presentation of these hallmarks was positively associated with aging (Figure 4B). Vascular cell senescence has been studied extensively across different tissues^78,79^, and emerging literature indicates that endothelial cell senescence is a prominent feature in the aging brain^53,80^. Microglial cell senescence has also been observed in the brain both in aging and disease, especially in the white matter^43,81^. Our unbiased pairwise gene coexpression analysis delineated unique gene combinations characteristic of white matter subpopulations of endothelial cells (Submodule 1.3) and microglia (Submodule 1.6). These submodules exhibit significant spatial and age-related correlation, and also contain key senescence-related genes (Figures 5C-E, Table S9).

Blood-brain barrier (BBB) integrity is maintained and modified by complex interactions between endothelial cells, pericytes, and resident immune cells, such as microglia. Senescent endothelial and microglial cells have the potential to disrupt complex interactions in the BBB, leading to BBB leakage and vulnerability^80^. Notably, Submodule 1.3, which is enriched in endothelial genes, includes MHC I antigens, while Submodule 1.6, which is enriched in microglia genes, includes MHC II antigens (Table S5). MHC antigen presentation has previously been identified as a means by which senescent cells are marked for immune cell clearance^82^. Recruitment of such immune cells from the periphery represents a potential mechanism by which senescent cells impact BBB function. A previous study in mice found that brain endothelial cells present MHC-I antigens to recruit CD8+ T cells to the BBB in the white matter of the spinal cord, contributing to BBB breakdown^83^. Further, mice with a liver-specific MHC-II deficiency showed impaired clearance of senescent hepatocytes by CD4+ T cells^84^. This suggests that alteration of antigen presentation may allow senescent endothelial and microglial cells to increase BBB vulnerability, contributing to age-related decline in dlPFC function.

Mature astrocytes have been shown to re-enter the cell cycle in response to tissue damage in the form of stroke, inflammation, and neurodegenerative disease pathology^85^. In our study, *AQP4* expression significantly increased in layers 4 and 5 of the dlPFC (Figure 2A and 2C). Both our spatial and snRNA-seq data identified an increase in astrocyte fraction with age (Figures 3A-E). This increase in astrocyte abundance is accompanied by changes in astrocyte gene expression programs. In our snRNA-seq dataset, nearly half (221 out of 452) of the identified differentially expressed genes were changing in astrocytes (Table S4). Among these aging markers were known senescence hallmarks, such as an increase in *HIF1a* (Figure 2F) and a decrease in *LMNB1* (Table S4) in aged astrocytes. Loss of LMNB1 has previously been documented as a hallmark of astrocyte senescence in the aged mouse hippocampus^86^.

We identified spatial astrocyte expression programs, submodules 5.3 and 13.1, that were strongly enriched with age (Figure 5F-K) and contain several senescence markers (Table S9). Additionally, many SASP factors expressed in astrocytes in our snRNA-seq dataset (Figure S6) were predicted to increase with age in our ST dataset, such as *AGT*, *BMP7*, *EFEMP1*, *IGFBP7*, *TGFB2,* and *VEGFA* (Figure 4F). Previous publications hypothesized a dual role of the SASP, with half of senescent cells adopting a pro-inflammatory, pro-fibrotic SASP profile, and the other half adopting a pro-growth, pro-tissue repair SASP profile^33,39^. Each of these mentioned SASP factors are growth factors, indicating a potential protective role senescent astrocytes play in the aging dlPFC. Additionally, *TGFB2*^72^, *BMP7*^72^ and *IGFBP7*^87^ secretion has been shown to promote senescence in surrounding cells in a paracrine dependent manner. Thus, the SASP profile of aged astrocytes of the dlPFC potentially promotes both tissue regeneration and senescence induction, highlighting the beneficial and deleterious roles senescent cells play in the body^24^.

Adult neurogenesis has not been reported outside the hippocampus or subventricular zone^88^. Despite this, mounting evidence suggests neurons can develop a senescence-like phenotype, though few studies indicate that neurons in healthy aged tissue express senescence markers^28^. We observed a significant number of excitatory neurons positive for senescence hallmarks, such as Fridman Senescence Up, p53 activated targets, and the DDR (Figure 4A). We also observed a subpopulation of excitatory neurons that expressed high levels of cell cycle arrest markers *CDKN2D* and *SATB1* (Figure 4C-D). *CDKN2D* expression was the main senescence marker found in neurofibrillary tangle-containing neurons in the dlPFC^41^. However, we unexpectedly found that the number of excitatory neurons positive for senescence hallmarks decreased with age (Figure 4B). More work is needed to determine whether the susceptibility of excitatory neuron subpopulations to develop senescence hallmarks drives the loss of excitatory neurons we observed in the aging dlPFC.

Our integrated spatial and snRNA-seq highlights multiple cell types in the human dlPFC that show both age-related changes and expression of subsets of senescence associated genes. These observed senescence phenotypes potentially serve different functions in the aging brain, as senescent cells play diverse physiological roles^16–23^. Current senolytic therapies such as D + Q act on senescent cells indirectly by disrupting senescent cell anti-apoptotic pathways^73,74^. However, such drugs act on senescent cells indiscriminately, resulting in the loss of pro-growth and pro-inflammatory senescent cells alike. Distinguishing between beneficial and harmful senescence and determining the molecular mechanisms driving the induction of distinct senescent phenotypes will help inform more targeted therapeutic strategies.

## Resource Availability

### Lead contact

Further information and requests for resources and reagents should be directed to and will be fulfilled by the lead contact, Hemali Phatnani.

### Materials availability

This study did not generate new unique reagents.

### Data and code availability

- Raw sequencing data will be made publicly available on the SenNet consortium portal upon publication: https://data.sennetconsortium.org/search?size=n_20_n&sort%5B0%5D%5Bfield%5D=last_modified_timestamp&sort%5B0%5D%5Bdirection%5D=desc
- Code used in this paper is publicly available on GitHub: https://github.com/JasonMares63/Columbia-TMC-DLFPC-Senescence/tree/main

## Acknowledgements

Research reported in this publication was supported by the National Institute of Aging of the NIH under award number U54AG076040. We are grateful to the San Diego TMC for allowing us to use their group’s senescence gene list. We thank the tissue donors whose generosity enabled this study.

## Author Contributions

Conceptualization, V.M., H.P.; methodology, N.S., J.M., A.D., V.M., H.P.; sequencing data production, S.G., N.B., O.C., K.K., S.K., B.F.; formal analysis, N.S., J.M., A.D.; writing, N.S., J.M., A.D., L.C., M.P., J.P., V.M., H.P.; writing – review and editing, all authors; resources, C.S., V.M., H.P.; data curation, J.M., A.D., C.M.; supervision, Y.S., H.P., V.M.; funding acquisition, C.S., Y.S., H.P., V.M.

## STAR Methods

### Resource Table

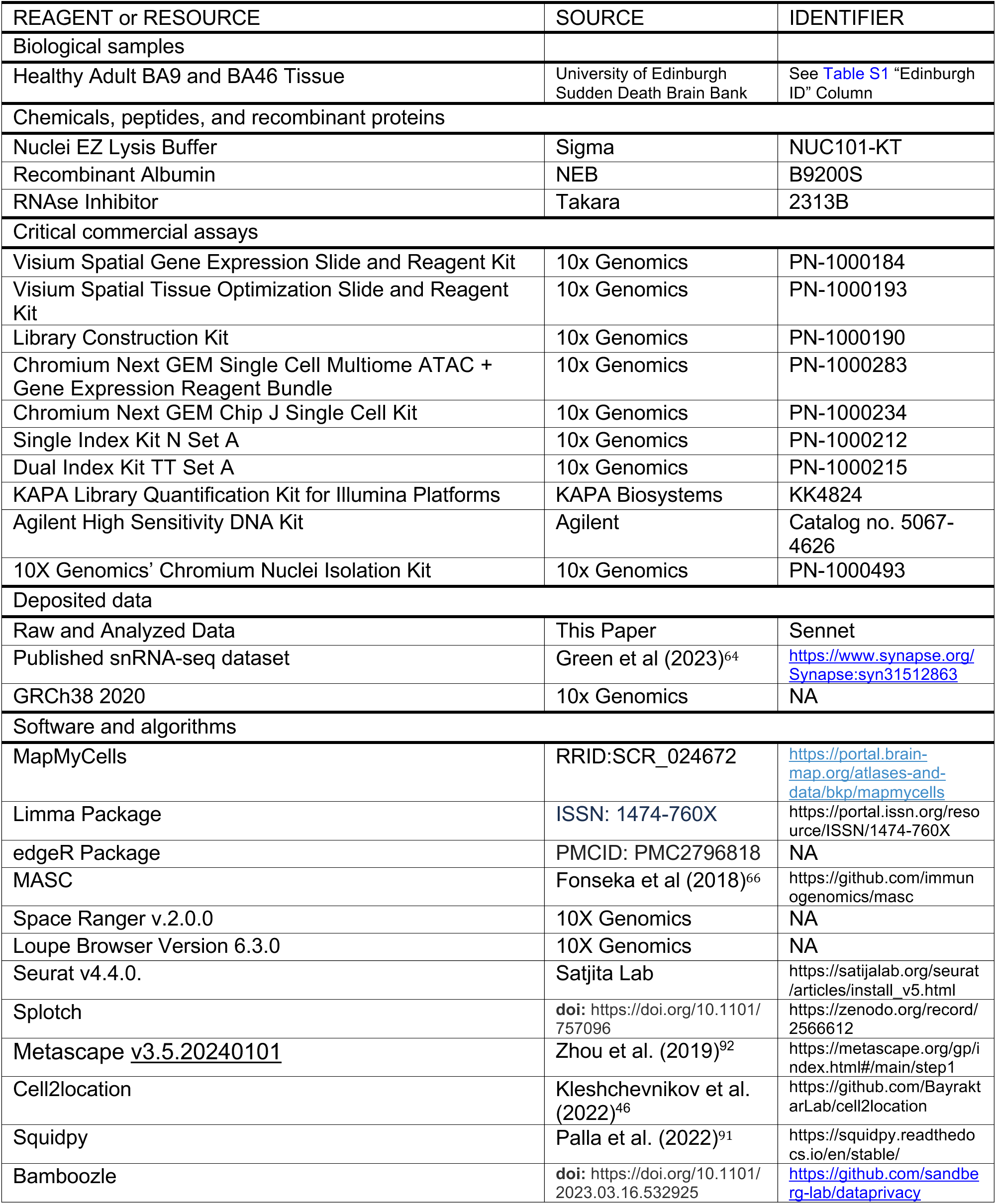

### TISSUE PREPARATION

#### Postmortem tissue

dlPFC tissue from sudden-death non-neurological control brains was acquired from the University of Edinburgh Sudden Death Brain Bank. Informed consent was acquired by the University of Edinburgh through its own institutional review board (IRB) protocol and samples were transferred to the Columbia University Tissue Mapping Center (CUSTMAP) in accordance with all applicable foreign, domestic, federal, state, and local laws and regulations for processing, sequencing and analysis. In depth subject metadata can be found in Table S1.

#### Tissue sectioning

Region BA9 or BA46 of the dlPFC was identified at autopsy, blocked out, and flash frozen on nitrogen vapors^89^. Frozen dlPFC tissue blocks were oriented perpendicular to the pial surface and trimmed to create sampling regions approximately 7 mm x 7 mm in size spanning the full gray matter from the pial surface to the white matter. Tissue blocks were required to have a balance of both gray and white matter to proceed with downstream experiments. Trimmed blocks were embedded in Tissue Plus OCT Compound (Fisher Healthcare, catalog no. 4585) and cryosectioned at -16°C.

For Visium ST, sections of 10 µm thickness cut perpendicular to the pial surface were collected onto prechilled Visium Spatial Gene Expression Slides (10x Genomics, catalog no. 1000185). Four technical replicates for each subject were collected across two Visium slides to minimize technical batch effects. Technical replicates on the same Visium slide are approximately 30-50 µm apart. Where necessary due to tissue sectioning artifacts, additional technical replicates were collected and processed on a later date.

For snRNA-seq, 5-10 sections of 50 µm thickness were cut from the same tissue block in the exact same plane that sections were cut for Visium ST experiments. This was to ensure we captured the same balance of cell types and cortical layers across both techniques for each subject. We then proceeded to the nuclei isolation protocol.

### SINGLE NUCLEI RNA SEQUENCING (snRNA-seq)

#### Nuclei Isolation protocol

Nuclei were isolated from frozen post-mortem dlPFC samples using a protocol previously published by Chatila et al. (2023)^90^. To profile nuclei from the same representative plane used in our Visium experiments, approximately 5-10 50 µm cryosections of post-mortem tissue were collected into pre-chilled 2 mL Eppendorf tubes. On ice, each sample received 2 mL of Nuclei EZ Lysis Buffer (Sigma; NUC101-KT) and was transferred to dounce tissue grinders (DWK Life Sciences, 357538). Samples were homogenized using ∼20-25 strokes with the loose pestle followed by ∼20-25 strokes with the tight pestle. The homogenized samples were transferred to a clean, pre-chilled 15 mL Falcon tube, where they received an additional 2 mL of Nuclei EZ Lysis Buffer. After pipette mixing, samples were incubated on ice for 5 minutes. Samples were centrifuged at 4C for 5 minutes at 500g. After removing the supernatants, nuclei pellets were resuspended in 4 mL of Nuclei EZ Lysis Buffer and incubated on ice for 5 minutes. Following a second centrifugation at 4C for 5 minutes at 500g, the supernatants were removed, and the nuclei pellets were resuspended in 1 mL of Nuclei Suspension Buffer (NSB: PBS, 0.01% BSA (NEB; B9200S), 0.1% RNAse inhibitor (Takara; 2313B)). The samples were centrifuged once more at 4C for 5 minutes at 500g. The nuclei pellets were resuspended in NSB, filtered through a 35 µm strainer, and counted. Each nuclei suspension was diluted for a targeted nuclei recovery of 5,000 nuclei per sample. Single nucleus RNA/ATAC sequencing was performed according to the 10X Genomics’ Chromium Next GEM Single Cell Multiome ATAC + Gene Expression User Guide (CG000338 Rev F).

A subset of nuclei from 3 subjects were isolated according to the 10X Genomics’ Chromium Nuclei Isolation Kit Use Guide (PN-1000493, CG000505, Rev. A). Metadata from one donor processed for snRNA-seq was incomplete and was therefore excluded from analyses assessing age-related trends in expression, but was included for cell-type annotation, cell-type deconvolution, and cell-type associated submodule formation (Table S1).

#### Sequencing

Size distribution and concentration of GEX libraries were assessed on the Fragment Analyzer and quantified using Picogreen and qPCR with the Universal KAPA Library Quantification Kit (catalog no. 07960255001). Libraries were pooled, loaded at 0.55 nM and sequenced on a NovaSeq 6000 System using the Illumina; NovaSeq 6000 S4 Reagent v1.5 Kit (200 Cycles) (catalog no. 20028313) kit. GEX libraries were sequenced using the following recipe: read 1: 28 reads, i7 index read: 10 cycles, i5 index read: 10 cycles, read 2: 90 cycles. Average sequencing depth for each GEX sample was approximately 143 x 10^6^ reads per library.

#### Quality check of transcriptomic data

Barcoded processing, gene counting and aggregation were made using the Cell Ranger software v2.0.0. All the 10X runs for each human sample were initially filtered with an nUMI cutoff of >500 and then nuclei with less than 5% mitochondrial gene contamination were retained. Next, the mitochondrial genes were also removed from the count matrices. DoubletFinder v3 was initially used to identify doublets, we leveraged these classifications during the clustering step to remove clusters with a high proportion of doublets and then any additional remaining doublets. Data was processed through the Seurat SCT workflow, including normalizing data, identifying variable features, and scaling. This was followed by running PCA, finding clusters, and projecting data onto UMAP space. Nuclei were manually annotated into seven broad classes: excitatory neurons, inhibitory neurons, oligodendrocytes, oligodendrocyte precursor cells (Opcs), astrocytes, microglia, vascular cells. We used marker genes: SNAP25 (for neurons), SLC17A7 (for excitatory neurons), GAD1 and GAD2 (for inhibitory neurons), MOG and MAG (for oligodendrocytes), PDGFRA (for Opcs), AQP4 and FGFRR3 (for astrocytes), TMEM119 and AIF1 (for microglia), and CLDN5 and PDGFRB (for vascular cells). After identifying broad classes, we split the data into these seven classes, and re-ran the SCT workflow to identify further contaminations in the data. A total of 53,007 nuclei passed quality control filtering, with mean detection of 1,897 genes per nucleus and 2,305 nuclei per sample (Table S1).

#### Mapped annotations of transcriptomic data

For further cell subtype annotation, we mapped our nuclei to the aged prefrontal cortex dataset released by Green et al. (2023)^64^. This comprehensive dataset includes 1.64 million single-nucleus RNA-seq profiles from 424 aging individuals. We used the Allen Institute cell type mapping algorithm, MapMyCells, (<u>RRID: SCR_024672</u>), a fast web-based tool. In addition, we ran this algorithm five times on our entire dataset and then used majority-vote from the five runs to select a final consensus annotation.

The output from the mapping algorithm annotated several nuclei into small subtype populations, as well as cell types that are not widely expected in the cortex. For these small clusters in glial cells, we merged clusters with less than 400 nuclei together. For the neurons, we merged pairs of clusters with less than 400 nuclei together that showed overlap in our latter run of Levain clustering. For cell types that were not expected in cortex, these were labeled as “Miscellaneous”. Lastly, we re-named these mapped clusters using their broad class name followed by their rank based on the number of nuclei found in our dataset. There are a total number of 37 mapped clusters, not including miscellaneous. We include a table that converts the cluster names we used in our paper with those used in the original Green et al. (2023)^64^ data (Table S11).

#### Single nucleus differential expression analysis

DEG analysis for the single nucleus RNA sequencing data was performed with the voom function from the limma package (ISSN: 1474-760X). Data was filtered, normalized, and aggregated using the edgeR package (PMCID: PMC2796818). We filtered for genes that were expressed in at least 30 nuclei and present in at least 4 samples. Afterwards, we performed linear modeling with limma where age is treated as a continuous variable. The voom function incorporates the mean-variance relationship in RNA-seq count data by assigning precision weights in order to assume normally distributed log-CPM values. All models included an intercept term along with a sex control variable. P-values were corrected for multiple testing using the Benjamini-Hochberg method. A summary of aging markers by cell type at the broad class and mapped subcluster level can be found in Table S4.

### Mixed association enrichment analysis

For associations between two categorical variables, we used MASC^66^, a tool for mixed effects <u>m</u>odeling of <u>a</u>ssociations of <u>s</u>ingle <u>c</u>ells. In this modeling, we used sex and batch as the fixed and random effects, respectively. For both MASC analyses in this paper, results for pericytes were not significant and confidence intervals were large relative to those for other subclusters. As a result, these latter results were not shown in figures.

### Single Nucleus Gene Module Scoring

Module scores were computed using Seurat’s *AddModuleScores* function. Scores are calculated by subtracting the average expression levels of gene sets at the single cell level with the average expression of a control feature set. This calculation is specifically done by, first, binning all genes (gene set and control set) based on average expression and then finding differences between the query genes and randomly selected control features. For calculating module scores for the snRNA-seq data of the 27 annotated gene lists, we maintain the same 2,400 control feature set when running *AddModule Scores*.

For each senescence hallmark, we calculated a threshold to determine whether nuclei could be classified as a senescence cell. The threshold for each hallmark was calculated following these steps (see figure below for further visualization of these steps):

1. For each gene in the hallmark that is also present in the snRNA-seq dataset, compute the percentage of nuclei expressing each marker with non-zero gene expression.
2. After determining each percentage value from (1), determine the median value of these percentages.
3. Calculate the threshold as the top *k*-th percentile of module scores across all nuclei where k is the median percentage from (2).

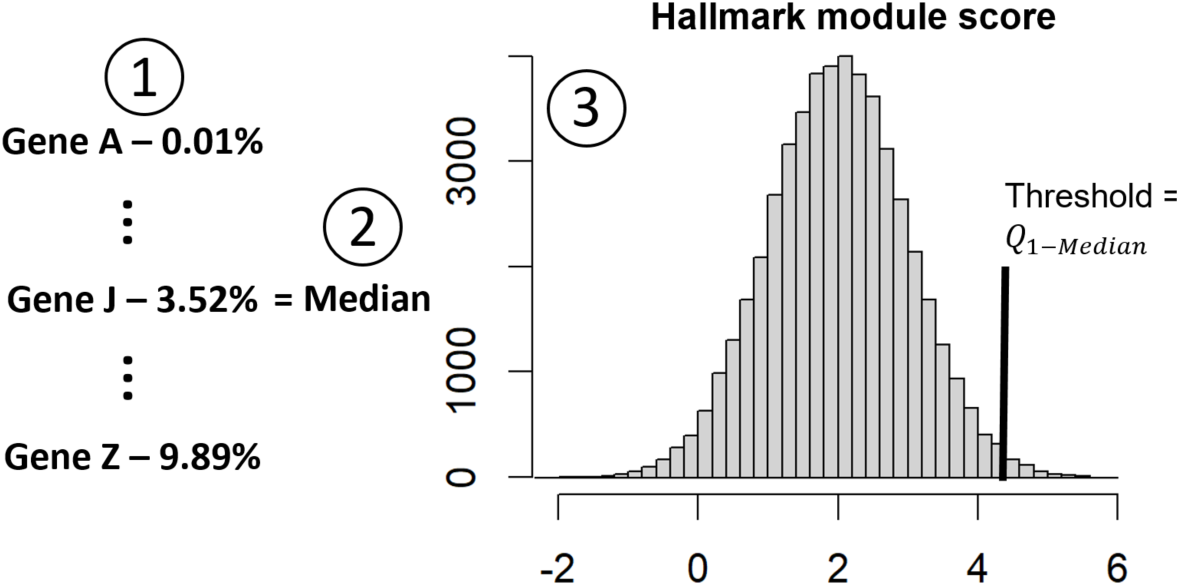

For each hallmark, we classify a nucleus as “positive” for that specific senescence hallmark if its module score is higher than the hallmark’s threshold (i.e. a threshold for each hallmark). We look into the level of overlap between nuclei labeled as “positive” across hallmarks through the usage of UpSet plots. These plots allow us to visualize the number of “positive” nuclei for each hallmark as well as which combination of hallmarks are showing the highest number of overlapping “positive” nuclei. For each hallmark and broad cell type, we test for differences in proportions of “positive” nuclei between samples from the young and middle/old age groups via a student’s t-test. Incorporating the SenNet consortium’s recommendation, we also identify nuclei that are positive for at least three senescent hallmarks, which we refer to as “3+ Hallmarks”. A summary of the number of positive nuclei for each broad class cell type and significance testing can be found in Table S10.

### VISIUM SPATIAL TRANSCRIPTOMICS (ST)

#### Tissue processing, imaging, and RNA capture for spatial transcriptomics

Visium spatially resolved gene expression data was generated according to the Visium Spatial Gene Expression User Guide (10x Genomics, CG000239 Rev F). Briefly, tissue sections collected onto Visium Spatial Gene Expression Slides (10x Genomics, catalog no. 1000185) were fixed in chilled methanol and an abbreviated hematoxylin and eosin protocol was used for histological staining. Brightfield RGB histological images were acquired using an EC Plan-Neofluar 10x/0.3 M27 objective on a Zeiss Axio Observer Z1 fitted with a Zeiss Axiocam 506 mono (Carl Zeiss Microscopy, Germany). Raw CZI images were stitched using Zen 2012 (blue edition) (Carl Zeiss Microscopy, Germany) and exported as JPEGs. Tissue sections were permeabilized for 12 minutes which was selected as the optimal time based on tissue permeabilization time course experiments conducted using the Visium tissue optimization protocol (10x Genomics, CG000238 Rev E). Reverse transcription was used to generate cDNA molecules incorporating spatial barcodes and unique molecular identifiers (UMIs). After second strand synthesis, cDNA was collected and amplified. The number of cDNA amplification cycles for each Visium array was identified using qPCR as described in the Visium Spatial Gene Expression User Guide.

#### cDNA library preparation

Visium libraries were manually prepared according to the Visium Spatial Gene Expression User Guide (10x Genomics, CG000239 Rev F) using unique Illumina-compatible PCR primers with Dual Index Kit TT Set A, 96 rxn (catalog no. 1000215, 10nt index). In addition to qPCR quantification using the KAPA Library Quantification Kit for Illumina Platforms (KAPA Biosystems, KK4824), a Qubit 3.0 fluorometer was used to measure the total yield of the final prepared libraries. Size distribution profiles of the final libraries were assessed using the Agilent 2100 Bioanalyzer system (Agilent High Sensitivity DNA Kit, catalog no. 5067-4626). Libraries that fell outside of the expected size range (mean library size of 450 bp with a range of 50 bp between libraries) and/or contained adapter dimer contaminations were flagged and excluded from the final pool. Library preparation was repeated for these samples and QCed as described.

#### Sequencing

Visium libraries were evaluated for size distribution on the Fragment Analyzer and quantified using Picogreen and qPCR with the Universal KAPA Library Quantification Kit (catalog no. 07960255001). Library pools were loaded at 0.7 nM concentration using the standard NovaSeq workflow as outlined in Illumina’s NovaSeq 6000 system guide (document #1000000019358 v12) and using the Illumina; NovaSeq 6000 S4 Reagent v1.5 Kit (200 Cycles)(catalog no. 20028313). Libraries were sequenced using the following recipe : read 1: 101 reads, i7 index read: 10 cycles, i5 index read: 10 cycles, read 2: 101 cycles. The average sequencing depth for each sample was approximately 239 x 10^6^ reads per Visium array and 59 x 10^3^ reads per spatial spot under tissue.

#### Sequence data preprocessing

Raw FASTQ files and histological images were processed using Space Ranger v.2.0.0, which uses a modified STAR v2.7.2a for genome alignment and performs barcode/UMI counting to generate feature-spot matrices. Reads were aligned to the GRCh38 2020-A human reference genome pre-built by 10x Genomics. Read alignment yielded an average of 1370 genes and 2882 UMIs per ST spot across our dataset before filtering of genes for Splotch analysis (Table S1).

Tissue arrays demonstrating extensive RNA degradation (with median genes detected per spot <700) were excluded from our analysis. Tissue sections with freeze fracture or other tissue sectioning artifacts that precluded cortical layer annotation of a majority of spots were also excluded from analysis. Where possible, these dropped replicates were replaced with additional technical replicates processed and sequenced on a later date. After array level QC, our dataset consists of 80 tissue sections from 8 young (33-44), 7 middle (45-55), and 7 old (56-74) subjects.

#### Cortical layer annotation

Following sequencing, each tissue section was manually annotated to capture transcriptomic changes occurring within unique cortical anatomical layers. All Visium spots under tissue sections were annotated with one of 7 anatomical annotation regions (AARs) consisting of the 6 cortical layers and the white matter. Annotation was performed using Loupe Browser (10x Genomics, Version 6.3.0) which enables interactive visualization of H&E images acquired as part of the Visium workflow with corresponding spot level gene expression data as registered by Space Ranger. Cortical layer annotations were assigned on the basis of cellular morphology in the portion of the histological image associated with each Visium spot. Cortical layers were identified as follows. The white matter was identified based on the lack of neuronal cell bodies and the linear arrangement of glial nuclei along neuronal processes. Cortical layer 6 was defined as the heterogeneous neuronal layer spanning from the white matter to the beginning of cortical layer 5, marked by large pyramidal neurons oriented perpendicularly to the pial surface. Cortical layer 4 was identified based on the presence of small, granular neurons adjacent to the large pyramidal neurons of cortical layer 5. Cortical layer 3 was labeled as the pyramidal cell layer between granular layers 2 and 4. Finally cortical layer 1 was identified based on the paucity of cells and lack of large neuronal soma. Spots that contained meninges were labeled as such and excluded from downstream analysis due to the distinct cellular composition of meninges as compared to cortical layer 1. Spots overlying tissue artifacts such as freeze fracture, folds, or tears were excluded from the analysis.

#### Spatial data integration and UMAP embedding

Visium ST data were integrated and projected into UMAP space using the anchor-based workflow of Seurat v4.4.0. Expression data were normalized and variable features selected for each Visium array using the NormalizeData and FindVariableFeatures methods, respectively. Integration features were selected across all arrays using SelectIntegrationFeatures, then scaling and PCA were performed over said features using ScaleData and RunPCA applied separately to each array. Due to the size of the dataset, reciprocal PCA (RPCA) with reference-based dataset anchors (FindIntegrationAnchors with 50 dimensions) was applied using all four arrays from one 50-year old male control donor (U54-HRA-010) as a reference. Finally, Visium data were integrated using IntegrateData, scaling was applied across the integrated dataset, and PCA was applied using default parameters, followed by further UMAP reduction of the first 50 PCs using min_dist=0.5 and n_neighbors=100 and all other parameters set to default values.

#### Splotch modeling of spatial gene expression data

Raw sequencing data was filtered to remove mitochondrially encoded genes, lncRNAs, pseudogenes, and genes expressed in less than 0.7% of spots. A summary of spot expression percentages for each gene in our dataset can be found in Table S3. Spots with fewer than 100 UMIs across remaining genes were discarded prior to Splotch analysis. Finally, spots without a valid annotation were discarded, and any spots without at least one immediate neighbor in the Visium grid were discarded to prevent singularities in the spatial autocorrelation component of the Splotch model. This resulted in expression data for 13,702 genes measured at 276,701 spatial spots across all patients with an average sequencing depth of 2,754 UMIs per spot.

Splotch employs a zero-inflated Poisson likelihood function (where the zero-inflation accounts for technical dropouts) to model the expression (*λ*) of each gene in each spot. This expression level is in turn modeled using a generalized linear model (GLM) with the following three components: the characteristic expression of the spot’s AAR (*β*), autocorrelation with the spot’s spatial neighbors (*ψ*), and spot-specific variation (*ε*). Furthermore, the characteristic expression rate (*β*) is hierarchically formulated to account for the experimental design, enabling the investigation of the effect of sample covariates on differential gene expression. By performing posterior inference on the model given the observed spatial expression data, we identified clinical group level trends in spatial gene expression (*β*) that best explains our observations. (*β*) is measured in log-scaled counts of gene expression per spot within an AAR of interest.

Compared to other computational methods for analysis of spatial transcriptomics data, Splotch i) enables quantification of expression differences between conditions and anatomical regions, ii) is designed to rigorously account for the uncertainty of low counts, and iii) analyzes multiple tissue sections simultaneously, stratifying patients based on clinical phenotype in order to quantify biological and technical variation.

### Spatial differential expression analysis

Differential expression analysis between conditions was performed by quantifying differences in the estimated posterior distributions (β) for young, middle and old subjects. This was accomplished using the Savage-Dickey density ratio to calculate a Bayes factor (BF) for a given comparison^57^ (e.g. expression of CDKN1A in cortical layer 1, Old vs Young). A BF >= 2 was the threshold used to identify differentially expressed genes between conditions. A BF>=30 was the threshold used to identify “top aging markers” for each AAR. Bayes factor tables summarizing differentially expressed genes with age can be found in Table S2.

### Spatio-temporal co-expression analysis

To study spatiotemporal and disease-dependent co-expression patterns in the human cortex, we consider all the mean posterior spot-level expression estimates (<u>*λ*</u>) from our Splotch model – a matrix with 276,701 rows (spots) and 13,702 columns (genes). First, we removed the effects of outlier spots by clipping <u>*λ*</u> values to the 99th percentile for each gene, then performed standard scaling on <u>*λ*</u> values (max value of 10) across spots separately for each donor to account for biological variation. Next, the gene-gene correlation matrix was calculated and hierarchical clustering (L1 norm and average linkage) applied to group genes of similar co-expression pattern across spots. The threshold for forming flat clusters was selected so that the main blocks on the diagonal belong to separate clusters. Any module with < 10 genes were discarded from further analysis. A summary of gene placement within our spatial coexpression modules can be found in Table S5.

For each coexpression module, cell type-associated submodules were derived using our donor-matched snRNA-seq dataset of 53,007 nuclei. The snRNA-seq data were transformed to transcripts per million (TPM), after which point any genes with a maximum expression of ≤ 10 TPM were discounted from submodule analysis due to insufficient expression. For all genes in a given ST-derived module, mean expression rates were calculated per cell type in the snRNA-seq data, and min-max scaled between 0 and 1 across the (k=37) cell types. Hierarchical clustering was then applied to these scaled average gene expression values, using the cosine distance and average linkage. The module genes were grouped into submodules by using the threshold 0.54*max(Z[:,2]), where Z is the linkage matrix, following the work of Maniatis et al. (2019)^57^. A summary of gene placement within our submodules can be found in Table S5.

Significance testing for differential module/submodule expression across age groups was accomplished by first calculating per-spot module scores, then averaging per-array and per-cortical layer to produce independent batch estimates and applying Welch’s t-test (with Benjamini-Hochberg FDR correction) between age groups (e.g., “Young” vs. “Old” in Layer_1). Per-spot module scores were calculated by first standard-scaling the expression of each module/submodule gene across all spots to remove the effects of within-module variation in basal gene expression, then averaging the expression of member genes within each spot.

### Gene ontology analysis of spatial modules

GO BP enrichment analysis of coexpression modules and cell type specific submodules was conducted using Metascape v3.5.20240101 (https://metascape.org/gp/index.html#/main/step1). Genes from modules or submodules used as “input list” while all other genes in our ST dataset were used as “background”. Enrichment was performed using the custom analysis tool, solely exploring enriched pathways from the “GO Biological Process” database, and with a p-value cutoff of 0.01. For the purpose of this analysis, only coexpression modules sized between 100 and 3000 genes were considered. A summary of GO BP terms enriched within each spatial coexpression module and submodule can be found in Table S6 and Table S7, respectively.

### Cell type deconvolution of Visium data

Per-spot estimates of the abundances of k=37 cellular subtypes were derived using Cell2location^65^. Cell2location accepts two data matrices: a (spots, genes) matrix of ST data and a (cells, genes) matrix of sc/snRNA-seq data. These data matrices were first subset to the set of genes detected in both modalities, then further subset to a set of marker genes “balanced” across the k=37 cell types (i.e., the same number of markers for each cell type) to enhance the differences between cellular subtypes.

Marker genes for deconvolution were derived from snRNA-seq data by first pseudo-bulking per-cell type and per-donor, then depth-normalizing to 1e4 counts per pseudo-sample, log-transforming, and applying scanpy’s rank_genes_groups method. Per-cell type, genes with >1.25 mean log-fold change (vs. other cell types) and p_adj<0.05 were retained, then each gene was assigned a dispersion score (variance-mean ratio within each cell type, averaged across cell types), and the top 1% of genes by dispersion were discarded following the methodology of Ma et al. (2018). Cellular subtype “Exc.8” was found to have the fewest (g=282) marker genes after filtering, and thus the superset of top g=282 marker genes per cell type (ranked by p_adj) were retained for deconvolution, resulting in 3,043 unique genes (Table S8).

Cell2location’s negative binomial (NB) regression model was trained on our full (53,007 nuclei, 3,034 genes) counts matrix (all donors) for 250 epochs with a batch size of 2,500 to produce reference signatures for all 37 cell types. Deconvolution of spatial data was performed per-donor with the aforementioned reference, treating individual Visium arrays as separate batches. The mean cells per spot was set to five, detection alpha set to 20 (default), and the model was trained for 30,000 epochs.

Significance testing for differential cell type abundance across age groups was accomplished by first normalizing spot composition vectors across cell types to sum to 1, then averaging per-array and per-cortical layer to produce independent batch estimates and applying the Wilcoxon rank sums test (with Benjamini-Hochberg FDR correction) between age groups (e.g., “Young” vs. “Old” in Layer_1).

### Gene set spatial enrichment analysis

Per-spot module scores were calculated for each of 30 gene sets (Table 9) using Seurat’s *AddModuleScores* function (*see Single Nucleus Gene Module scoring* methods for more details). For the control feature set, we randomly selected 2,400 genes not included in any of the 30 gene sets and used this same list across all gene sets.

Deciles were computed for each module across all n=276,701 spots in the dataset, and each spot was assigned a discrete label between 1 (d1) and 10 (d10) for each module based on the decile of the observed expression score. Separately for each age group and module, spatial enrichment scores for all pairs of deciles were calculated using Squidpy’s^91^ `nhood_enrich` function with n=1,000 permutations, treating individual Visium arrays as batches.

## Supplemental Information

**<u>Figures S1-S10:</u> In main document**

**<u>Table S1</u>: Donor metadata and QC metrics, related to figure 1**

In depth subject metadata sheet and basic ST/snRNA-seq subject by subject QC metrics

**<u>Table S2</u>: ST DEG analysis, related to figures 2**, **4 and 5**

Splotch 1 Bayes Factor Tables (Level 1 hierarchy, meaning Old vs Young, Middle vs Young, and Old vs Middle)

**<u>Table S3</u>: Spot level expression percentages, related to figures 2**, **4, and 5**

Spot expression percentage of each gene in our splotch run (e.g. CDKN1A expressed in 1.3% of spots)

**<u>Table S4</u>: snRNA-seq DEG analysis, related to figure 2**

snRNA-seq DEG analysis at the broad class level and subcluster level

**<u>Table S5</u>: Module and submodule gene lists, related to figure 5**

Gene placement in spatial coexpression modules and cell type specific submodules

**<u>Table S6</u>: GO BP pathways enriched in spatial modules, related to figure 5**

Gene ontology biological processes (GO BP) pathways enriched for each module (100-3000 gene modules only)

**<u>Table S7</u>: GO BP pathways enriched in spatial cell type submodules, related to figure 5**

Gene ontology biological processes (GO BP) pathways enriched for each submodule of interest

**<u>Table S8</u>: Cell2location markers, related to figure 3**

Marker genes used for cell2location deconvolution of mapped cluster cell types

**<u>Table S9</u>: Senescence hallmark gene lists and submodule annotation, related to figures 4**, **5, and 6**

Table summarizing all curated senescence gene sets used in our analysis

**<u>Table S10</u>: Broad class senescence module enrichment results, related to figure 4**

Table summarizing statistics from figure 4a

**<u>Table S11</u>: Subcluster labels, related to figure 3**

Conversion table between Green et al. (2023)^64^ subclusters and paper sub clusters nomenclature

