## Supplementary material for "Uncovering the Signatures of Cellular Senescence in the Human Dorsolateral Prefrontal Cortex": Figures S1-10

### Supplementary Figures

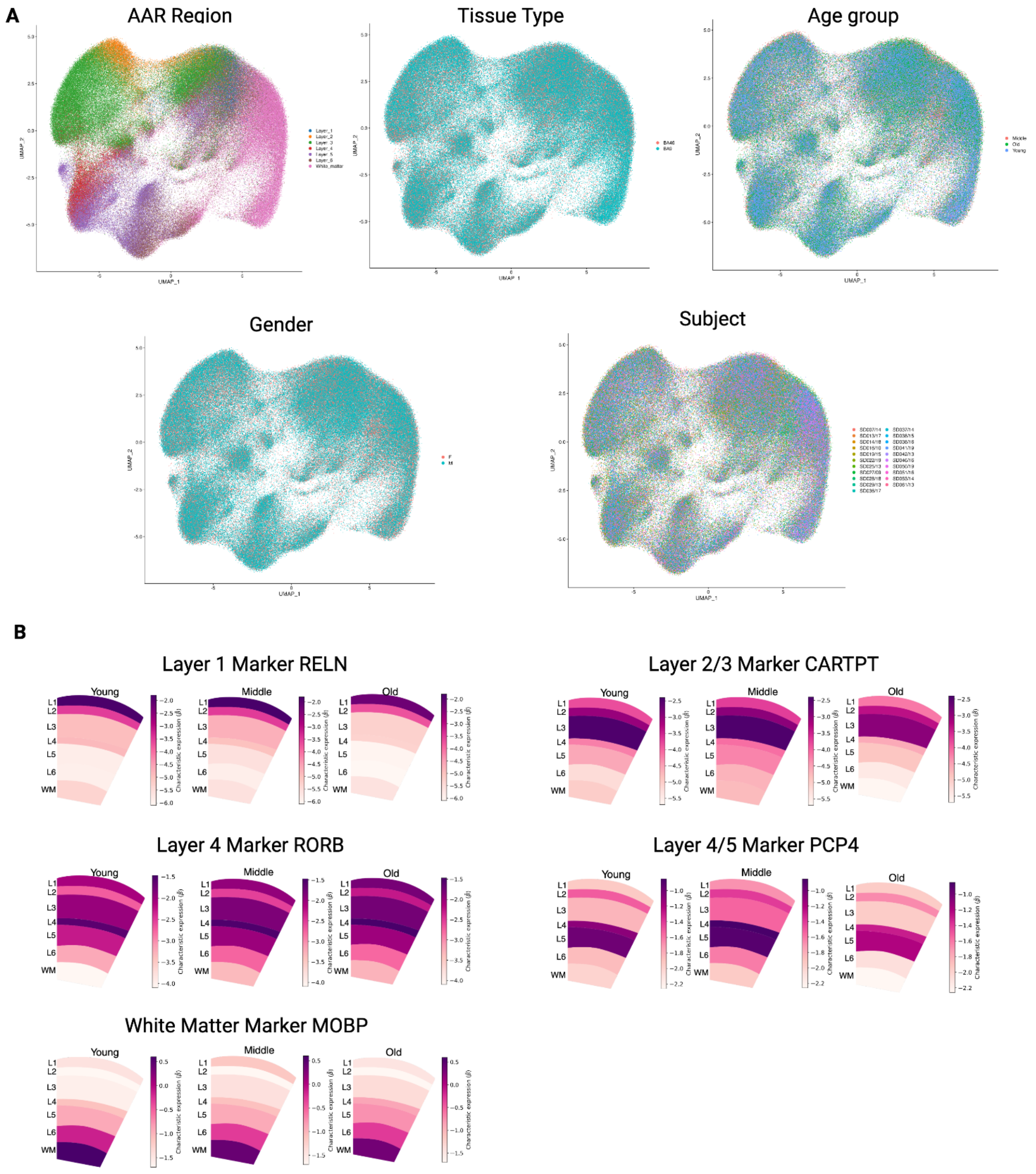


**Figure S1 Additional QC of ST dataset, related to figure S1**
**(A)** UMAP plots of Visium data, colored by AAR region, tissue type, age group, gender, and subject, respectively. **(B)** AAR level Splotch-predicted expression of key cortical layer markers within reference ST arrays from young, middle, and old age groups

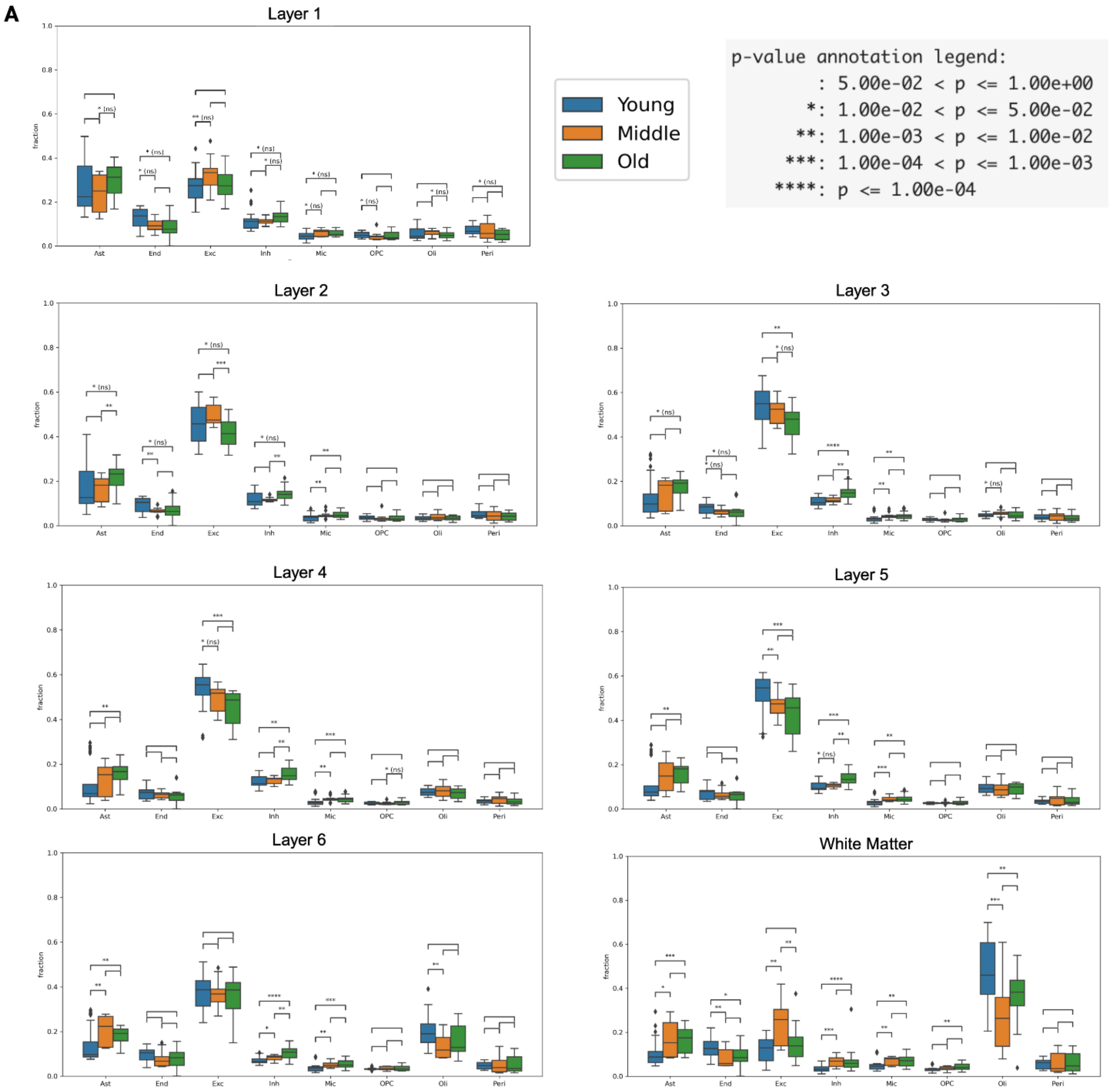


**Figure S2 Changes in broad class cell type composition by layer, related to figure 3**
**(A)** Broad class cell type deconvolution output for ST data shown across cortical layers and age groups. Pairwise differential testing results between age groups shown on top of box plots, along with reported p-values.

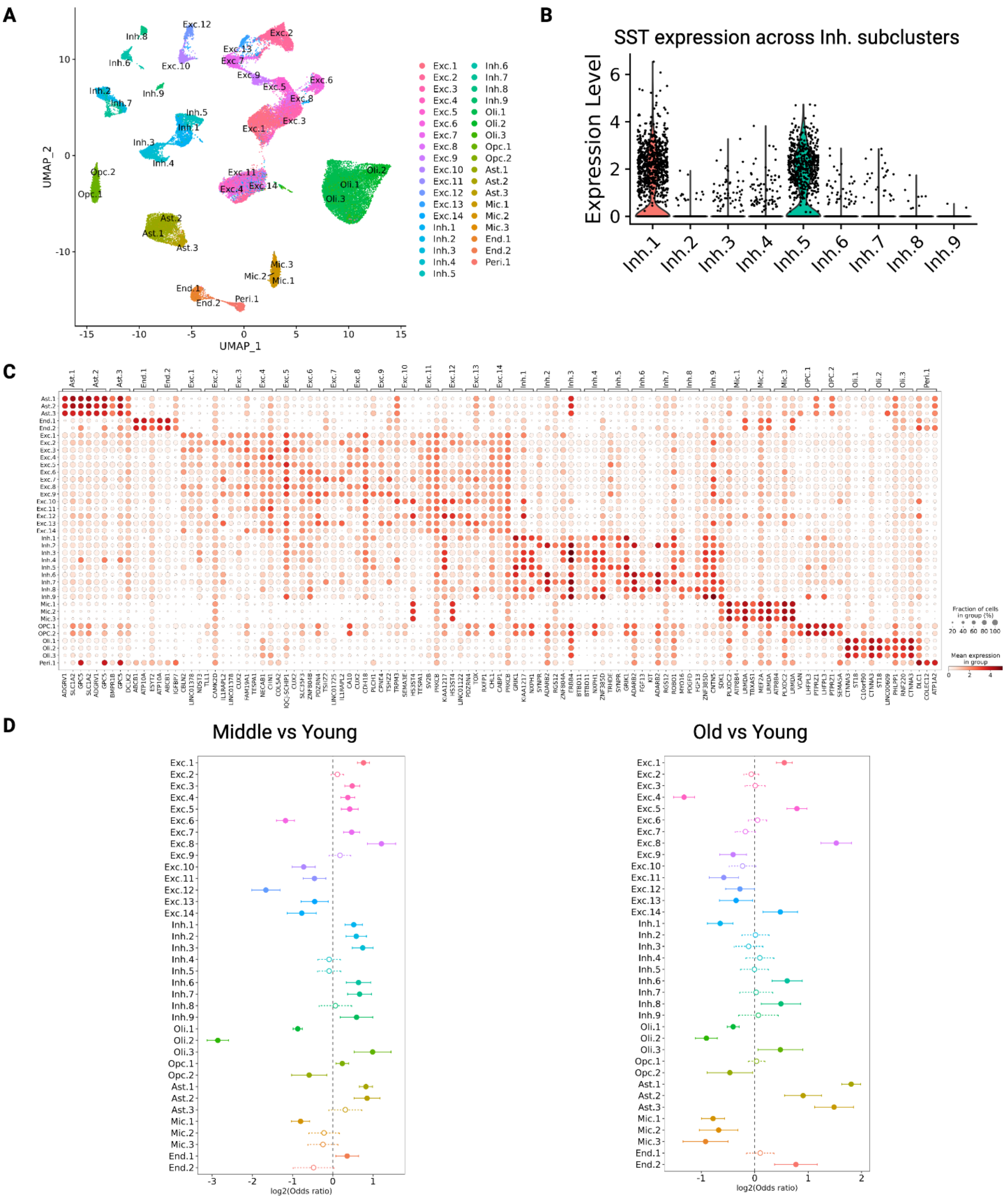


**Figure S3 Mapped subcluster markers and changes in composition, related to figure 3**
**(A)** UMAP plot of subclusters of the snRNA-seq data. (**B**) Expression level of SST across Inh. subclusters. **(C)** Summary of markers specific to cell type subclusters identified in our snRNA-seq dataset. A full list of markers can be found in Table S4. **(D)** MASC differential cell type proportion testing results between young and middle (left) and young and old (right) groups are shown. Points to the left (right) indicate increased (decreased) cell type enrichment, respectively, in the older group. Significant results (p<=.05) are indicated by solid, filled-in bubbles and solid confidence interval bars.

**A**

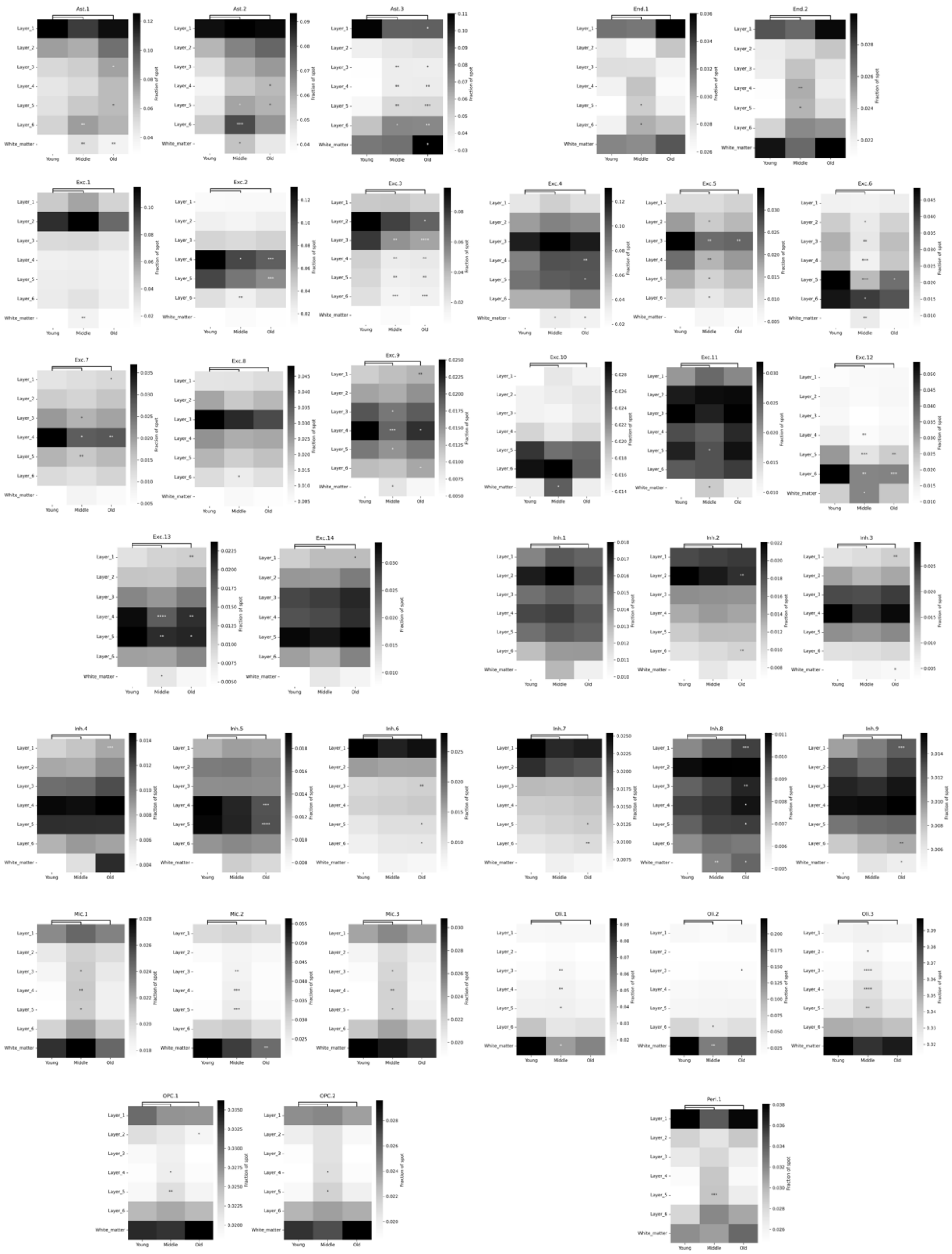

**Figure S4 Extended cell2location mapped subcluster results, related to figure 3**
**(A)** Heatmaps visualize abundance of all mapped clusters in ST data across cortical layers predicted by cell2location. Asterisks are added to highlight significant results from differential abundance testing between middle and old, versus young groups. For heatmaps, asterisks indicate level of significance: ****=p<1e-4***=p<1e-3 **=p<1e-2, *=p<.05.

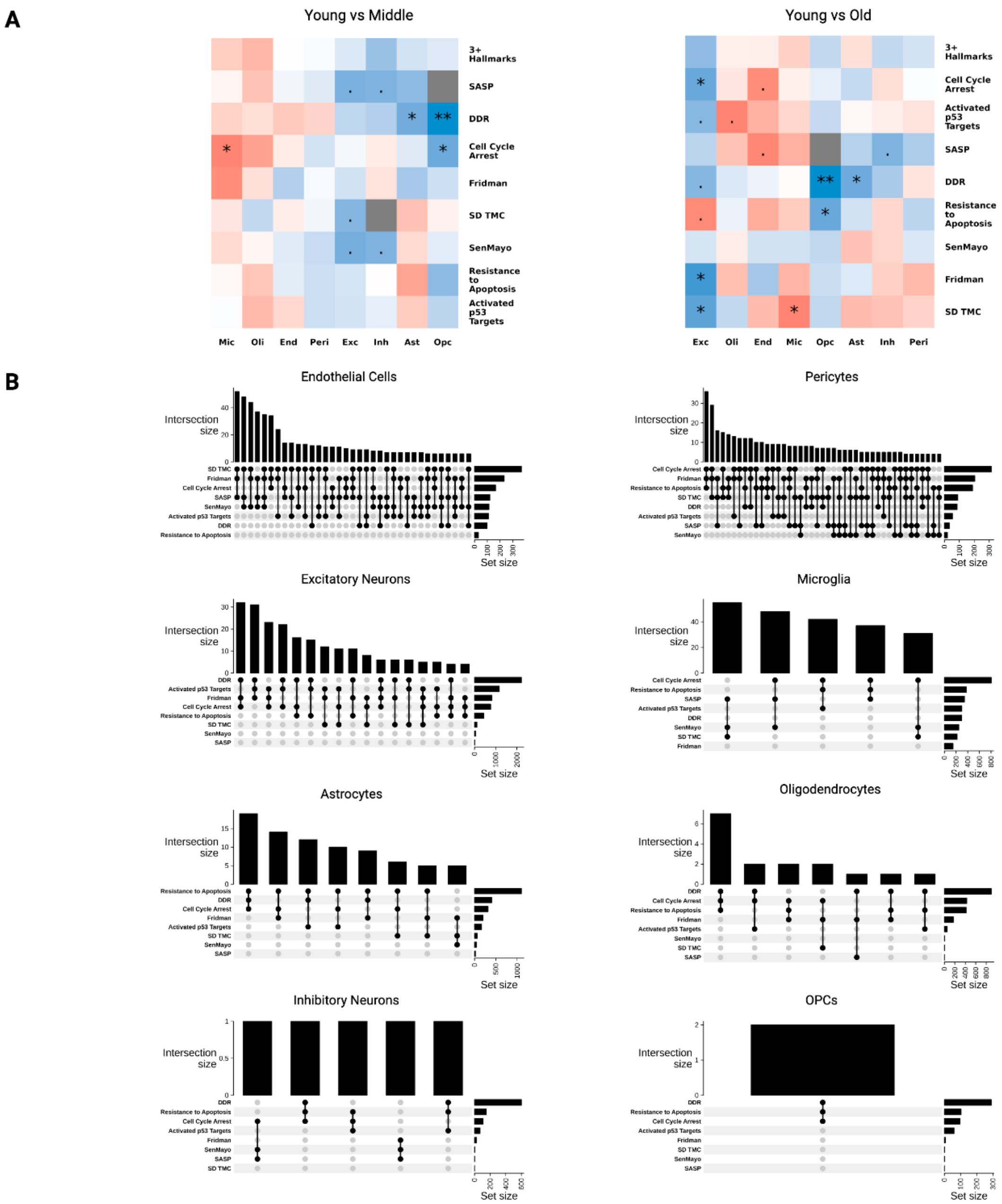


**Figure S5 Figure 4a/b extended results, related to figure 4**

**(A)** Heatmaps displaying enrichment of senescent-positive nuclei between young and middle/old (left/right) cohorts across senescence hallmarks. Cells represent test statistics from testing for difference of proportions via t-test. Nuclei that are positive for the "3+ Hallmarks" meta hallmark are nuclei that are positive for at least three senescence hallmarks. Red (blue) cells indicate an increased proportion of senescent nuclei in the old (young) cohort. Nominal p-values are annotated onto cells. **=p<1e-2, *=p<.05, .=p<=.10. **(B)** UpSet plots quantifying the overlap of nuclei positive across at least three senescence hallmarks by broad class cell type. For each UpSet plot, bar plots at the top indicate the number of overlapping nuclei while the bar plot to the right indicates the number of positive nuclei for each hallmark.

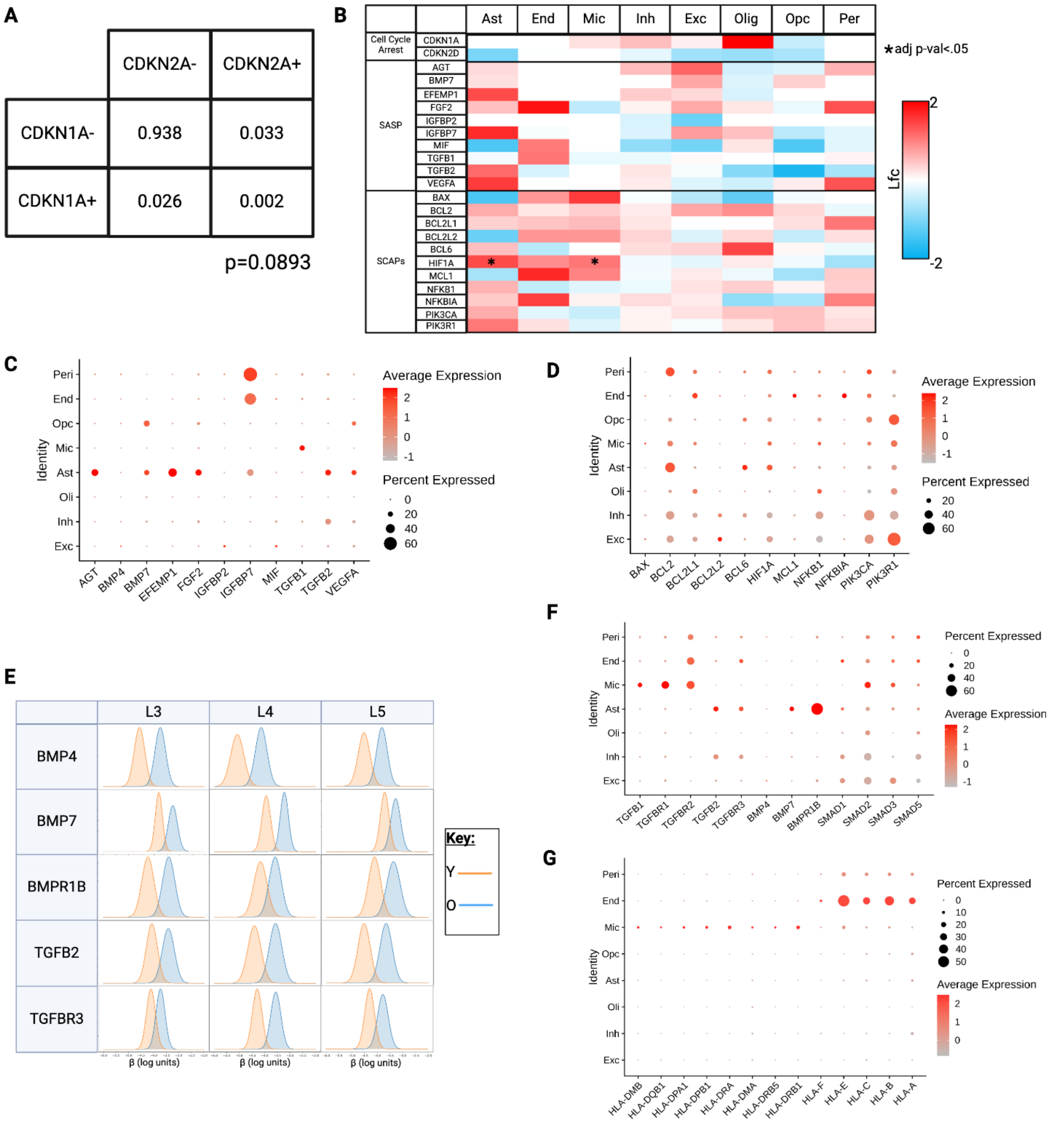


**Figure S6 Expression pattern of senescence genes and TGF-β family members, related to figure 4**

**(A)** Percentage of microglia positive/negative for CDKN1A/CDKN2A. Statistical testing failed to determine dependency of CDKN1A+ and CDKN2A+ microglial populations. **(B)** Heatmap displaying age-related changes in expression of senescence related genes in snRNA-seq dataset (complementary to Figure 4F). **(C-D)** Dot plots displaying average mRNA expression level of SASP and SCAPs members, respectively, listed in **(B)**. **(E)** Differential expression between young (Y) and old (O) subjects in various TGF-β family ligand and receptor members measured by ST. **(F)** Average expression levels of TGF-β family members transcripts across our snRNA-seq dataset. (**G**) Average expression levels of HLA family members transcripts across our snRNA-seq dataset.

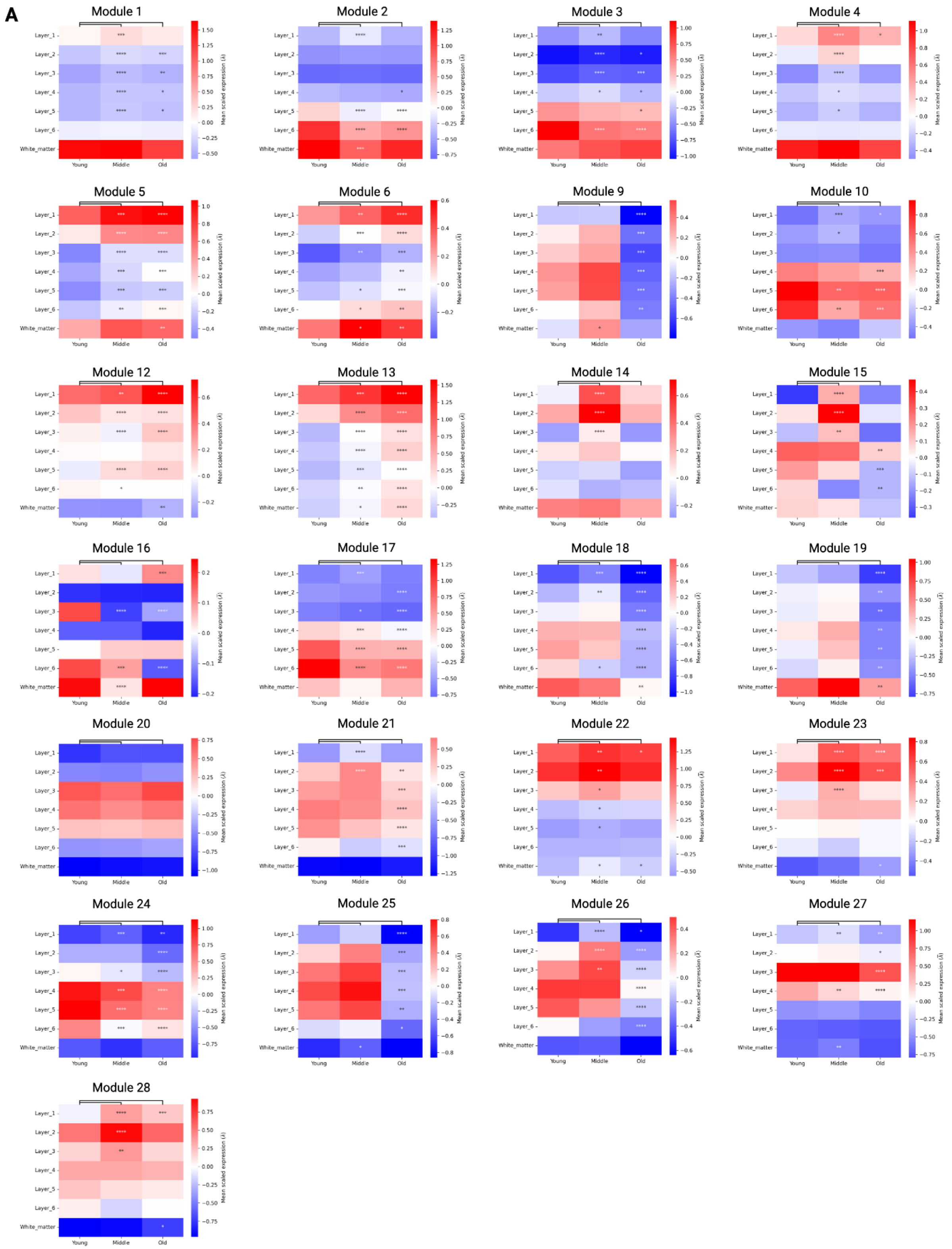


**Figure S7 pt1 Spatial module and cell-type specific submodule activities, related to figure 5**

**(A)** Average expression levels of each spatial module generated in Figure 5 by cortical layer and age groups, with differential testing results between old and middle versus young. Asterisks indicate level of significance: ****=p<1e-4***=p<1e-3 **=p<1e-2, *=p<.05.

**B**

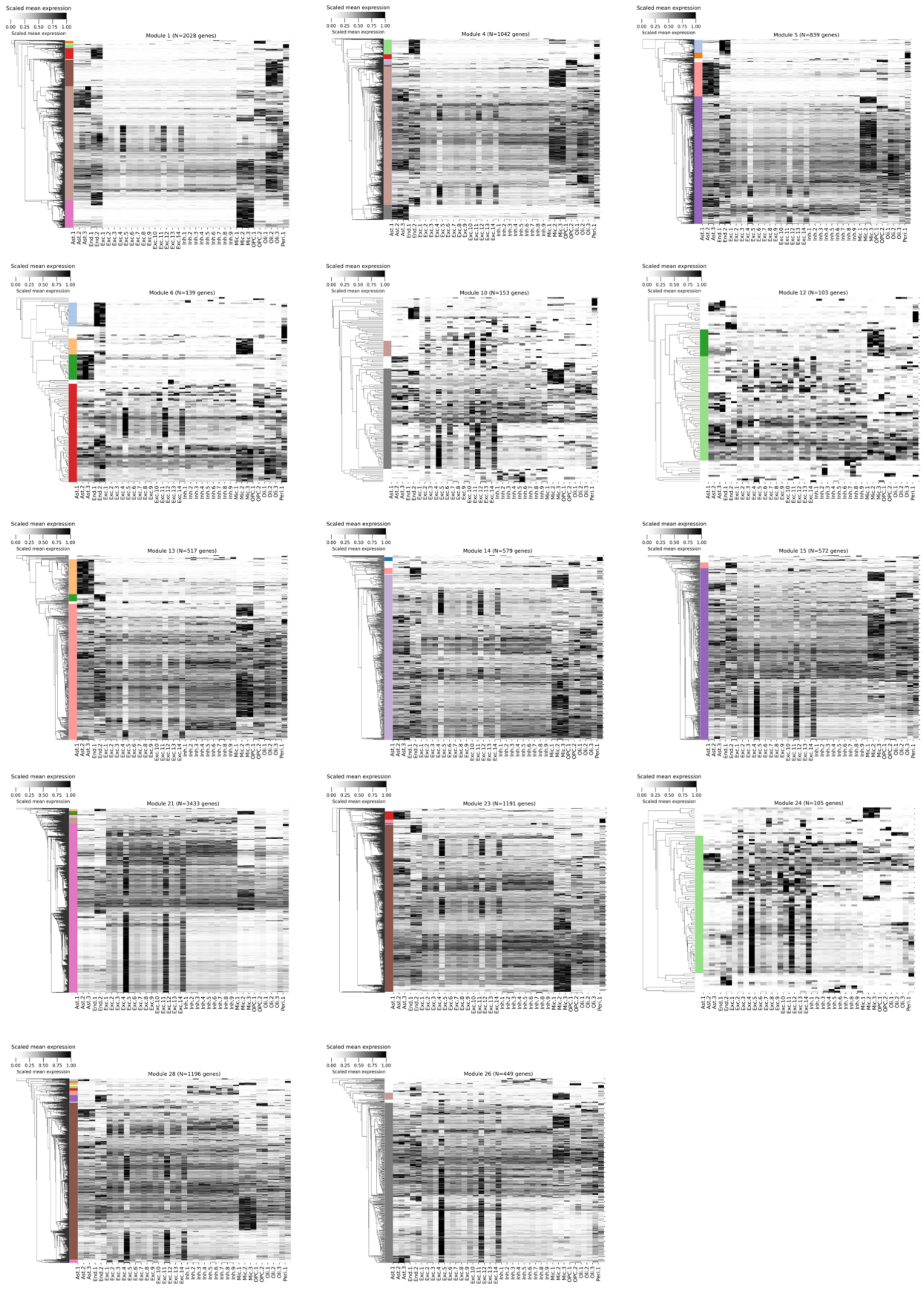


**Figure S7 pt2 Spatial module and cell-type specific submodule activities, related to figure 5**

**(B)** Heatmaps showcase expression patterns of genes in each spatial module (minimum 100 genes) generated in Figure 5 across cell types using donor matched snRNA-seq data.

c

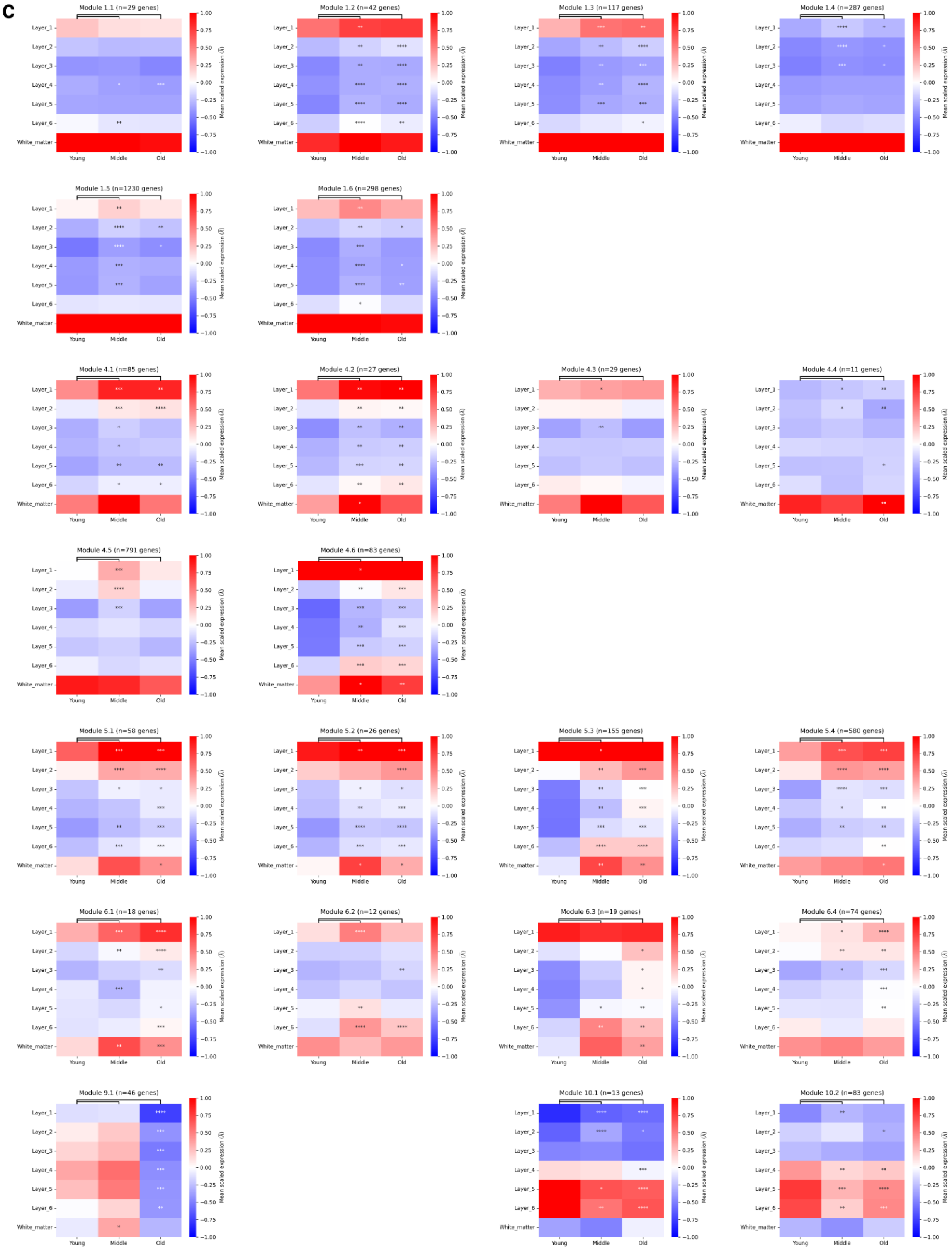
Module 1.1 (n=29 genes)
Module 1.2 (n=42 genes)
Module 1.3 (n=117 genes)
Module 1.4 (n=287 genes)
Module 1.5 (n=1230 genes)
Module 1.6 (n=298 genes)
Module 4.1 (n=85 genes)
Module 4.2 (n=27 genes)
Module 4.3 (n=29 genes)
Module 4.4 (n=11 genes)
Module 4.5 (n=791 genes)
Module 4.6 (n=83 genes)
Module 5.1 (n=58 genes)
Module 5.2 (n=26 genes)
Module 5.3 (n=155 genes)
Module 5.4 (n=580 genes)
Module 6.1 (n=18 genes)
Module 6.2 (n=12 genes)
Module 6.3 (n=19 genes)
Module 6.4 (n=74 genes)
Module 9.1 (n=46 genes)
Module 10.1 (n=13 genes)
Module 10.2 (n=83 genes)
Layer_1
Layer_2
Layer_3
Layer_4
Layer_5
Layer_6
White_matter
Young
Middle
Old
Mean scaled expression (λ̄)
1.00
0.75
0.50
0.25
0.00
−0.25
−0.50
−0.75
−1.00

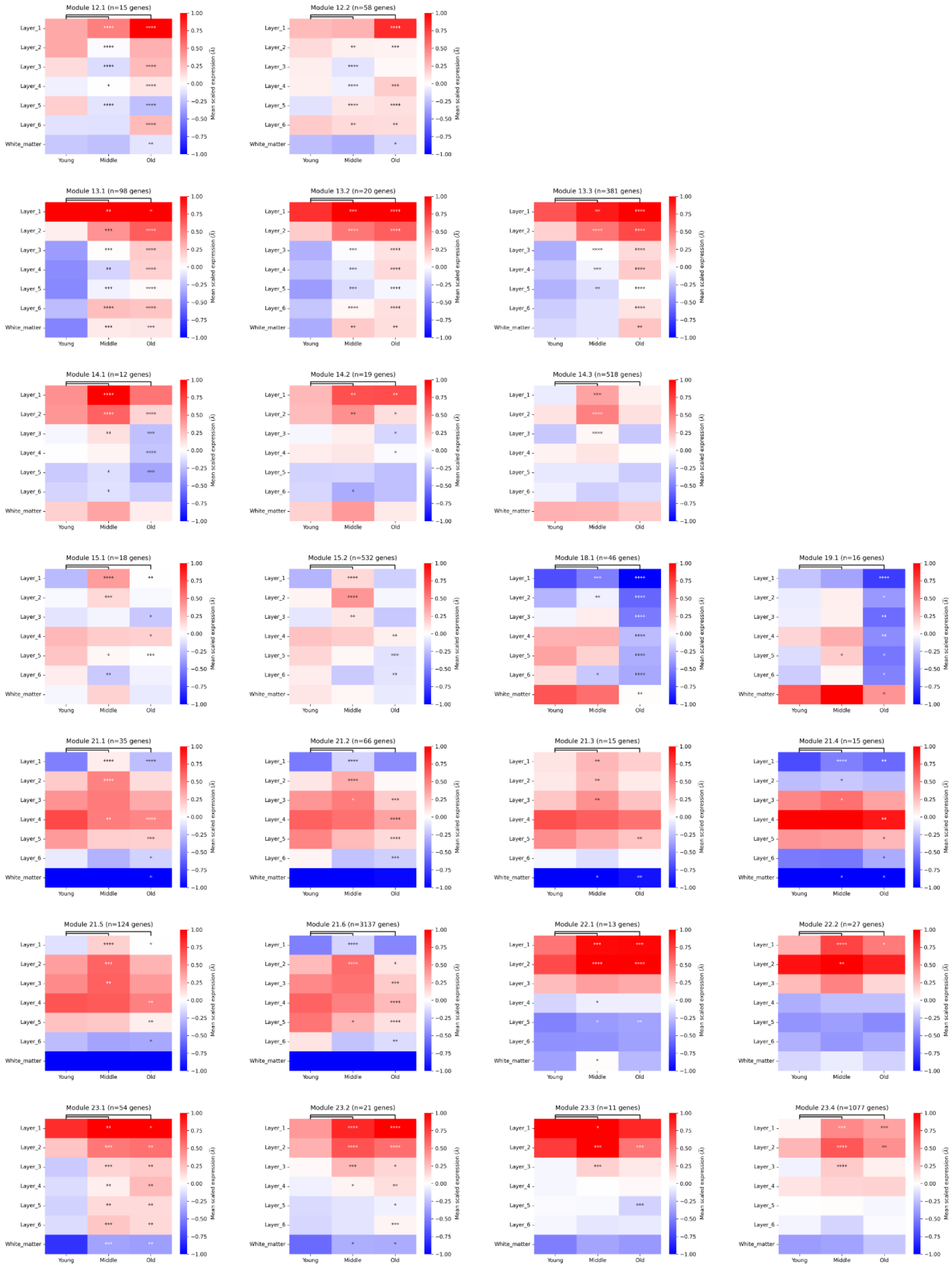

Module 12.1 (n=15 genes)
Module 12.2 (n=58 genes)
Module 13.1 (n=98 genes)
Module 13.2 (n=20 genes)
Module 13.3 (n=381 genes)
Module 14.1 (n=12 genes)
Module 14.2 (n=19 genes)
Module 14.3 (n=518 genes)
Module 15.1 (n=18 genes)
Module 15.2 (n=532 genes)
Module 18.1 (n=46 genes)
Module 19.1 (n=16 genes)
Module 21.1 (n=35 genes)
Module 21.2 (n=66 genes)
Module 21.3 (n=15 genes)
Module 21.4 (n=15 genes)
Module 21.5 (n=124 genes)
Module 21.6 (n=3137 genes)
Module 22.1 (n=13 genes)
Module 22.2 (n=27 genes)
Module 23.1 (n=54 genes)
Module 23.2 (n=21 genes)
Module 23.3 (n=11 genes)
Module 23.4 (n=1077 genes)
Layer_1
Layer_2
Layer_3
Layer_4
Layer_5
Layer_6
White_matter
Young
Middle
Old
Mean scaled expression (λ̄)
1.00
0.75
0.50
0.25
0.00
−0.25
−0.50
−0.75
−1.00

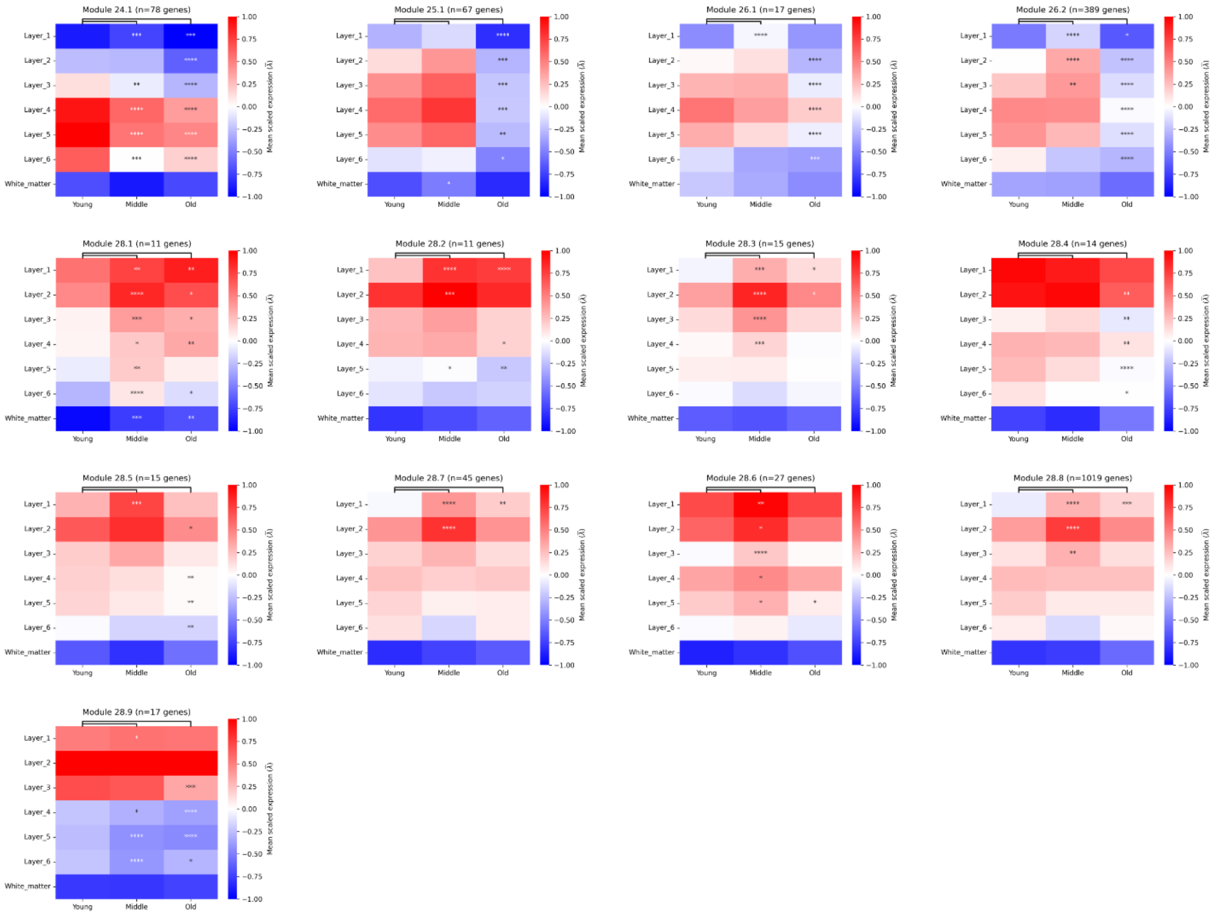


**D**

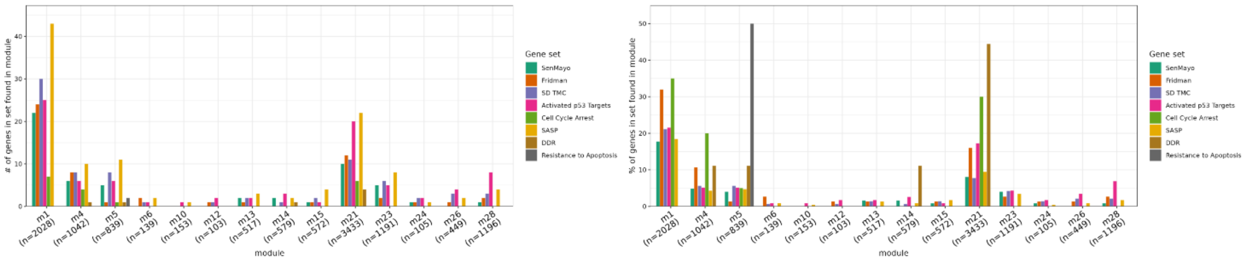


**Figure S7 pt3 Spatial module and cell-type specific submodule activities, related to figure 5**
**(C)** Average expression levels of each spatial cell type specific submodule generated in Figure 5 by cortical layer and age groups, with differential testing results between old and middle versus young. Asterisks indicate level of significance: ****=p<1e-4***=p<1e-3 **=p<1e-2, *=p<.05. (**D**) Bar plots displaying number (left) and percentage (right) of senescence hallmark genes from each curated list found in each spatiotemporal module generated in figure 5.

**A**

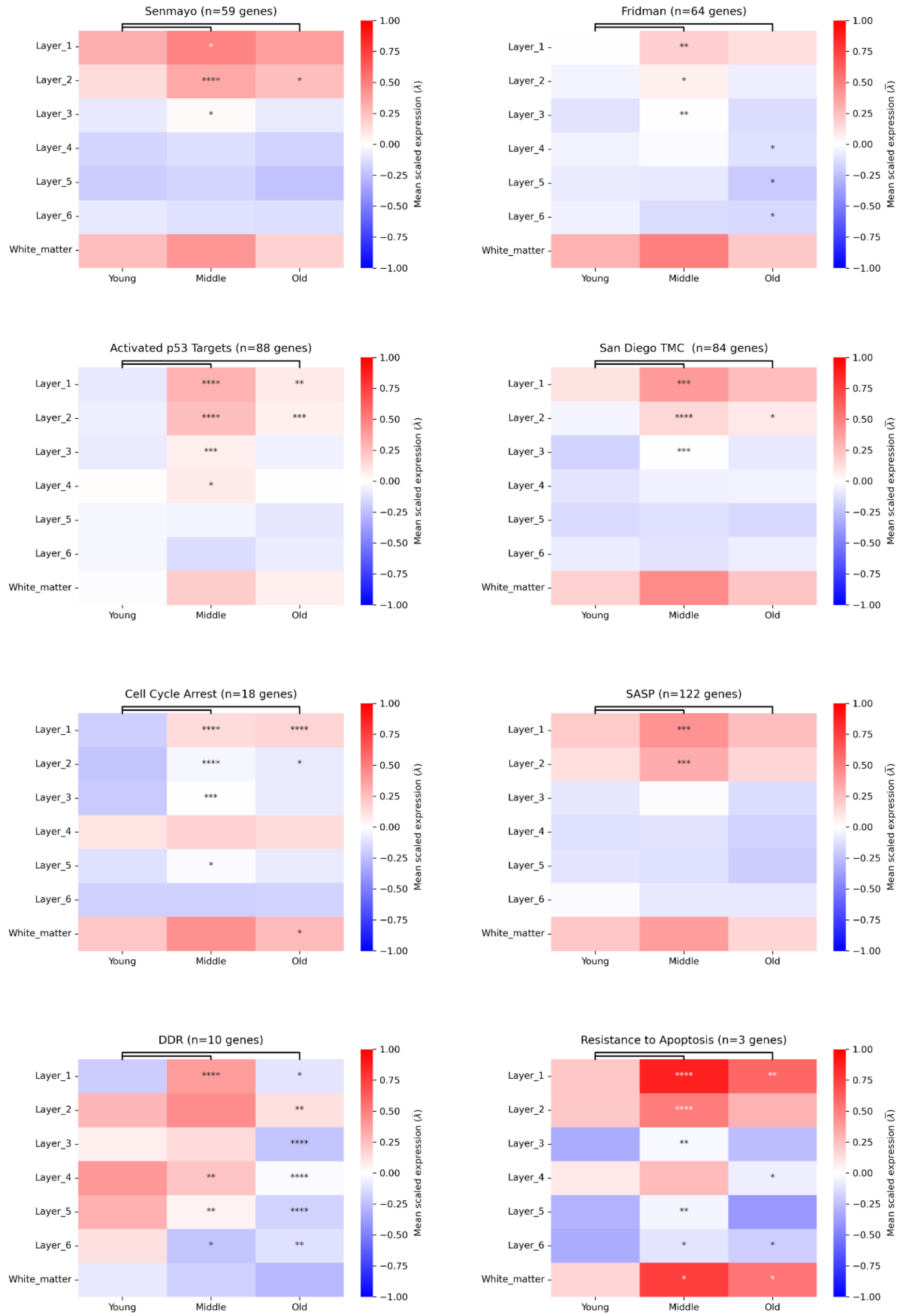


**Figure S8 Mean expression levels of senescence hallmarks measured by ST, related to figures 4, 5, and 6**

**(A)** Average expression levels of each curated senescence hallmark list by cortical layer and age groups, with differential testing results between old and middle versus young. Asterisks indicate level of significance: ****=p<1e-4***=p<1e-3 **=p<1e-2, *=p<.05.

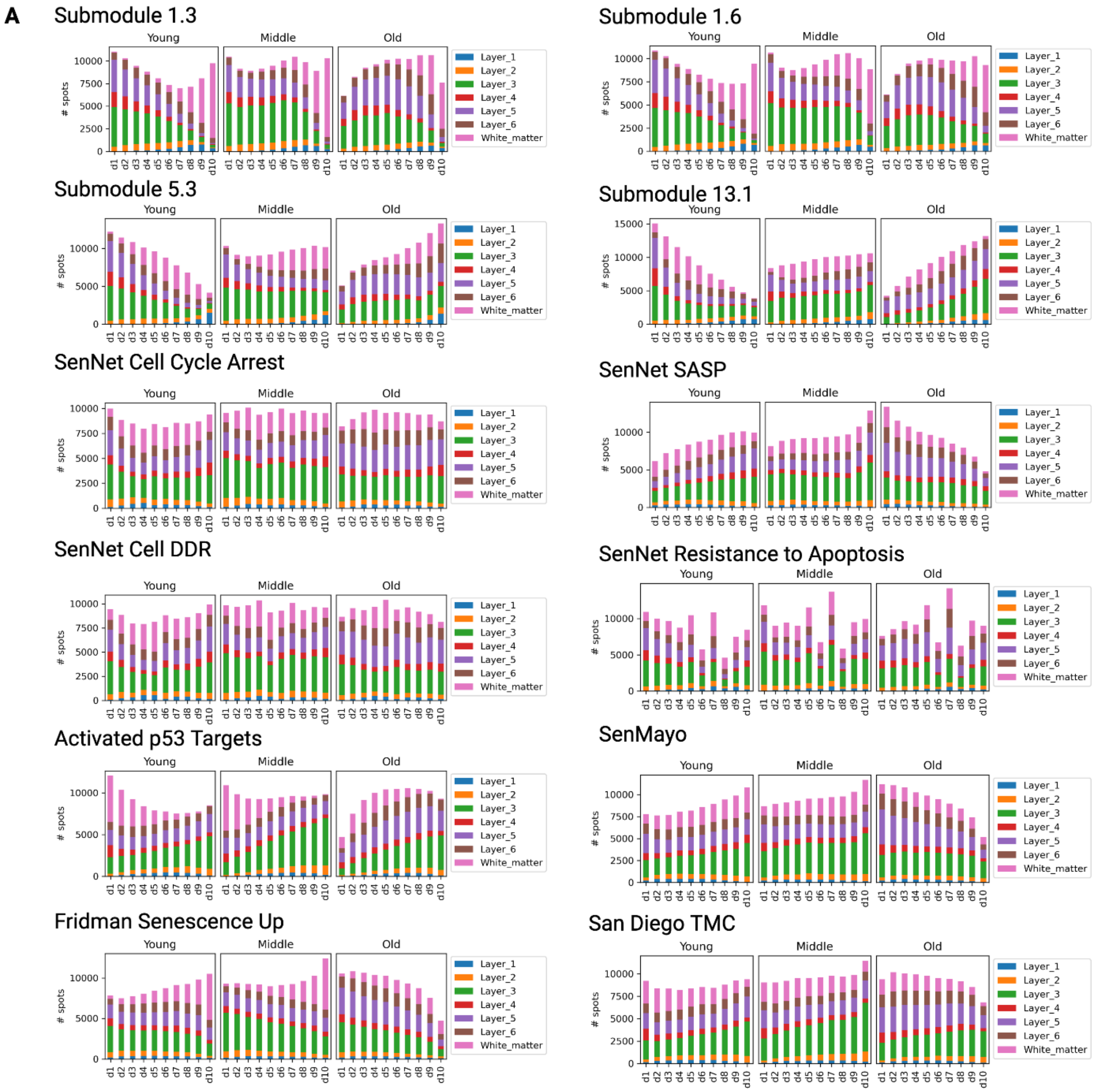


**Figure S9 Senescence module decile barplots, related to figure 6**

**(A)** Spatial transcriptomic spots were scored for expression of each unbiased spatial submodule of interest as well as known senescence modules from the literature. Module scores across ST spots were divided into deciles. We then determined the distribution of spots scored at each decile for young, middle, and old age groups, with stacked bar plots showing AAR locations of all spots separated by age and module expression.

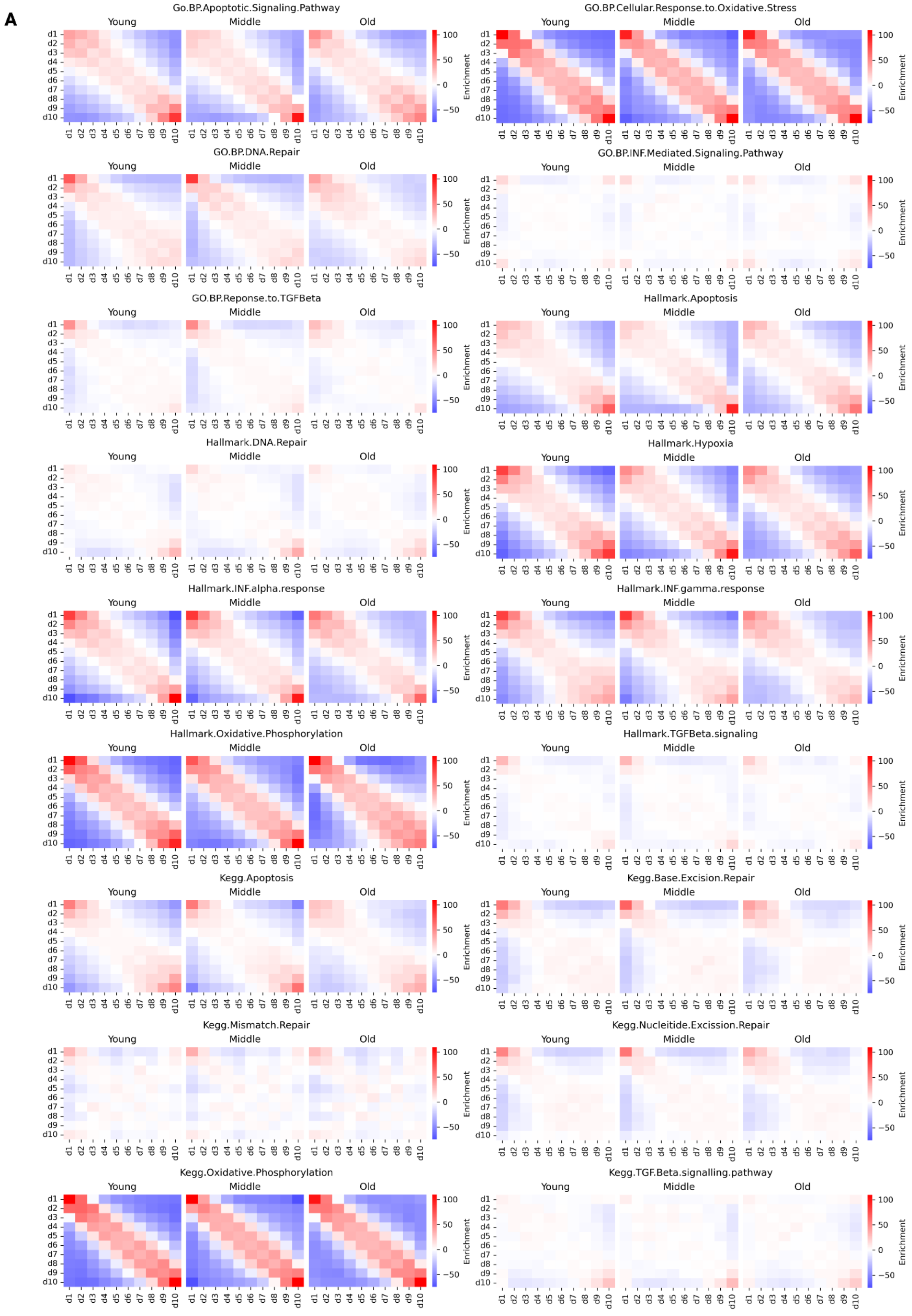
A
Go.BP.Apoptotic.Signaling.Pathway
GO.BP.Cellular.Response.to.Oxidative.Stress
GO.BP.DNA.Repair
GO.BP.INF.Mediated.Signaling.Pathway
GO.BP.Reponse.to.TGFBeta
Hallmark.Apoptosis
Hallmark.DNA.Repair
Hallmark.Hypoxia
Hallmark.INF.alpha.response
Hallmark.INF.gamma.response
Hallmark.Oxidative.Phosphorylation
Hallmark.TGFBeta.signaling
Kegg.Apoptosis
Kegg.Base.Excision.Repair
Kegg.Mismatch.Repair
Kegg.Nucleitide.Excission.Repair
Kegg.Oxidative.Phosphorylation
Kegg.TGF.Beta.signalling.pathway
Young
Middle
Old
Enrichment
100
50
0
−50
d1
d2
d3
d4
d5
d6
d7
d8
d9
d10

**B**

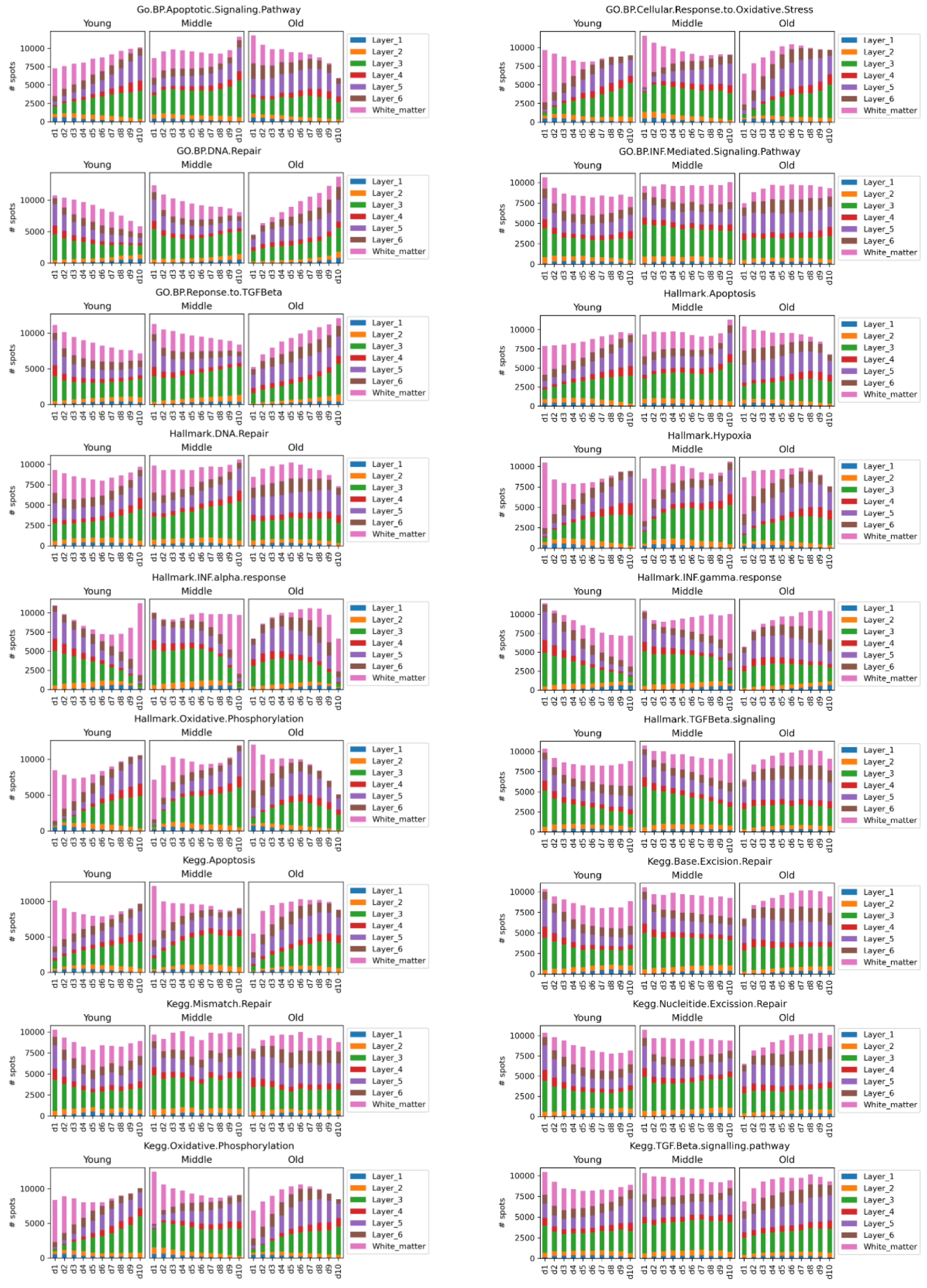


**Figure S10 Spatial distribution of public curated pathways with age, related to figure 6**

**(A-B)** Spatial transcriptomic spots were scored for expression of GO, Kegg, and Hallmark pathways with relevance to senescence (see Table S9 for full gene lists). Module scores across ST spots were divided into deciles. For each spatial submodule, we display heatmaps showing spatial enrichment statistics for each decile and quantified self enrichment (**A**). We then determined the distribution of spots scored at each decile for young, middle, and old age groups, with stacked bar plots showing AAR locations of all spots separated by age and module expression (**B**).
